# BIN1 recovers tauopathy-induced long-term memory deficits in mice and interacts with Tau through Thr^348^ phosphorylation

**DOI:** 10.1101/462317

**Authors:** Maxime Sartori, Tiago Mendes, Shruti Desai, Alessia Lasorsa, Adrien Herledan, Nicolas Malmanche, Petra Mäkinen, Mikael Marttinen, Idir Malki, Julien Chapuis, Amandine Flaig, Anaïs-Camille Vreulx, Philippe Amouyel, Florence Leroux, Benoit Déprez, François-Xavier Cantrelle, Damien Maréchal, Laurent Pradier, Mikko Hiltunen, Isabelle Landrieu, Devrim Kilinc, Yann Herault, Jocelyn Laporte, Jean-Charles Lambert

## Abstract

The bridging integrator 1 gene (*BIN1*) is a major genetic risk factor for Alzheimer’s disease (AD). In this report, we investigated how BIN1-dependent pathophysiological processes might be associated with Tau. We first generated a cohort of control and transgenic mice either overexpressing human MAPT (Tg*MAPT*) or both human MAPT and BIN1 (Tg*MAPT*;Tg*BIN1*), which we followed-up from 3 to 15 months. In Tg*MAPT*;Tg*BIN1* mice short-term memory deficits appeared earlier than in Tg*MAPT* mice; however – unlike Tg*MAPT* mice – Tg*MAPT*;Tg*BIN1* mice did not exhibit any long-term or spatial memory deficits for at least 15 months. After sacrifice of the cohort at 18 months, immunohistochemistry revealed that BIN1 overexpression prevents both Tau mislocalization and somatic inclusion in the hippocampus, where an increase in BIN1-Tau interaction was also observed. We then sought mechanisms controlling the BIN1-Tau interaction. We developed a high-content screening approach to characterize modulators of the BIN1-Tau interaction in an agnostic way (1,126 compounds targeting multiple pathways), and we identified – among others – an inhibitor of Calcineurin, a Ser/Thr phosphatase. We determined that Calcineurin dephosphorylates BIN1 on a Cyclin-dependent kinase phosphorylation site at T348, promoting the open conformation of the neuronal BIN1 isoform. Phosphorylation of this site increases the availability of the BIN1 SH3 domain for Tau interaction, as demonstrated by nuclear magnetic resonance experiments and in primary neurons. Finally, we observed that the levels of the neuronal BIN1 isoform were decreased in AD brains, whereas phospho-BIN1(T348):BIN1 ratio was increased, suggesting a compensatory mechanism. In conclusion, our data support the idea that BIN1 modulates the AD risk through an intricate regulation of its interaction with Tau. Alteration in BIN1 expression or activity may disrupt this regulatory balance with Tau and have direct effects on learning and memory.

## Introduction

Alzheimer’s disease (AD) is the most common neurodegenerative disorder and is clinically characterized among others by memory deficits affecting first short term and then long term and spatial memory. AD constitutes a major public, medical, societal, and economic issue worldwide, with 35.6 million people suffering from the disease and a forecast of 106 million in 2050.^1^ Responding effectively to this AD crisis necessitates a better understanding of this disease in order to improve diagnosis and therapy.

AD is characterized by two main types of brain lesions: (i) amyloid plaques, resulting from the extracellular accumulation of amyloid beta (Aβ) peptides; (ii) neurofibrillar degeneration, due to the intracellular aggregation of abnormally hyperphosphorylated Tau proteins. This latter aggregation is associated with an abnormal localization of Tau from the axonal compartment to the somato-dendritic compartment.^2^

The discovery of mutations in the *APP*, *PS1* and *PS2* genes (coding for amyloid precursor protein, APP, and presenilins 1 and 2), responsible for early-onset, autosomal-dominant forms of AD, has placed Aβ oligomer production at the center of the pathophysiological process.^3^ A better understanding of the genetic component of the common, complex forms of AD, which is exceptionally high among multifactorial aging-related diseases,^4^ is required to decipher the pathophysiological processes of AD. Genome-wide association studies (GWAS) allowed for the identification of more than 30 loci associated with the late-onset forms of AD,^5-8^ including the bridging integrator 1 gene (*BIN1*). A part of these genes pointed out a potential failure in Aβ clearance, leading to more insidious Aβ accumulation in the brain.^8, 9^ On the other hand, it is only recently that AD genetic risk factors have been also associated with Tau pathology, following the development of systematic screenings in *Drosophila* which allowed for the identification of genetic modifiers by assessing eye roughness and eye size as readouts of Tau neurotoxicity ^10-12^ and their associations with endophenotypes related to Tau.^10, 13, 14^ Such observations are of high importance since, contrary to amyloid plaques, neurofibrillary tangles (NFTs) are well correlated with cognitive impairment both in humans ^15^ and in animal models.^16^

Among the genes described to genetically interact with human Tau transgene in *Drosophila*, *BIN1* was further described to directly interact with the Tau protein by NMR spectroscopy using recombinant proteins, *in vitro* glutathion *S*-transferase (GST) pull-down from HEK293 lysates, as well as reciprocal co-immunoprecipitation from mouse brain synaptosome homogenates.^17^ In addition, a genome-wide significant functional risk variant in the vicinity of *BIN1* locus has been associated with Tau loads (but not with Aβ loads) in AD brains.^10^

The *BIN1* gene codes for Amphiphysin 2, also called BIN1, a ubiquitously expressed protein involved in membrane remodeling. BIN1 comprises a N-BAR domain involved in membrane curvature sensing, an SH3 domain that binds to proline-rich motifs present in a number of proteins including itself, and a clathrin- and AP2-binding domain (CLAP) specific of the neuronal isoform 1.^18^ In the central nervous system (CNS), BIN1 is mostly found in the axon initial segment, at the nodes of Ranvier,^19^ and at the synapse,^20, 21^ and was also associated with myelinated axons and oligodendrocytes in the white and grey matter.^22^ However, little is known about its function in the CNS. We recently described the consequences of increased human BIN1 expression in the mouse brain, which exhibits early alterations in the neuronal tract between the entorhinal cortex and the dentate gyrus of the hippocampus, leading to impaired novel object recognition and aging-related changes.^21^ Altogether, BIN1 overexpression affects the aging brain and induces neurodegeneration.^21^

Little is also known about BIN1 in the context of AD. Several teams evaluated potential links between AD and BIN1 and determined: (i) BIN1 may regulate BACE1 intracellular trafficking through multiple mechanisms and subsequently alter Aβ peptide production;^23^ (ii) BIN1 may have a role in plasma membrane remodeling during myelination, which is known to be affected in AD;^22, 24^ (iii) BIN1 may participate in the neuron-to-neuron propagation of Tau prion strains;^25^ and (iv) BIN1 may directly interact with Tau and interfere with Tau neurotoxicity *via* unknown mechanisms.^10, 26^ Strong efforts are thus still needed to determine how BIN1 is involved in the pathophysiological processes of AD. In this study, we assessed for the first time the impact of human *BIN1* overexpression in a mouse model of tauopathy and further dissected the interaction between Tau and BIN1 at the molecular and cellular levels using multidisciplinary approaches.

## Results

### *BIN1* overexpression modulates hTau phenotypes in short- and long-term memory

Although a genetic interaction between Bin1 and *MAPT* has been shown in *Drosophila* and the corresponding proteins have been described to physically interact,^17, 26^ the impact of BIN1 expression levels on cognitive function has not yet been investigated in a mammalian tauopathy model. For this purpose, we crossed the hTau mouse, a tauopathy model that overexpresses human *MAPT* (but does not express endogenous murine *Mapt* ^27^) with the Tg*BIN1* mouse that overexpresses human *BIN1* under the control of its own promoter and recapitulates the tissue-specific expression of different BIN1 isoforms.^21^ Briefly, generation of mice were obtained on C57BL/6J genetic background by crossing *Mapt*^+/-^;Tg*MAPT/0* ^28^ and *Mapt*^+/-^;Tg*BIN1/0* ^21^ to obtain *Mapt*^+/-^ as control littermates, *Mapt^-/-^*;Tg*MAPT/0* (noted here hTau) as the tauopathy model,^27^ and, finally, *Mapt^-/-^;*Tg*MAPT*/0;Tg*BIN1/0* as the double transgenic model (noted here hTau;Tg*BIN1*). Notably, in the Tg*BIN1* mouse, brain *Mapt* expression is similar to that observed in the WT mouse (Fig. S1).

To assess if *BIN1* overexpression affected the short-term, non-spatial memory deficit in the hTau mice, a novel object recognition (NOR) task was performed longitudinally at 3, 6, 9, 12, and 15 months. *MAPT* overexpression induced short-term memory deficits in males and females from 9 months on, characterized by their inability to discriminate between familiar and novel objects (Fig. 1A). Strikingly, hTau;Tg*BIN1* mice displayed short-term memory deficits earlier than hTau mice, by 3 months, both in males and females. Notably, *Mapt* heterozygous deletion alone had no impact on this task and Tg*BIN1* males present NOR deficits only starting from 6 months.^21^ There was no place or object preference, regardless of genotype or sex (Fig. S2). In conclusion, hTau phenotypes in the NOR task appeared at an earlier age upon *BIN1* overexpression.

**Figure 1.**
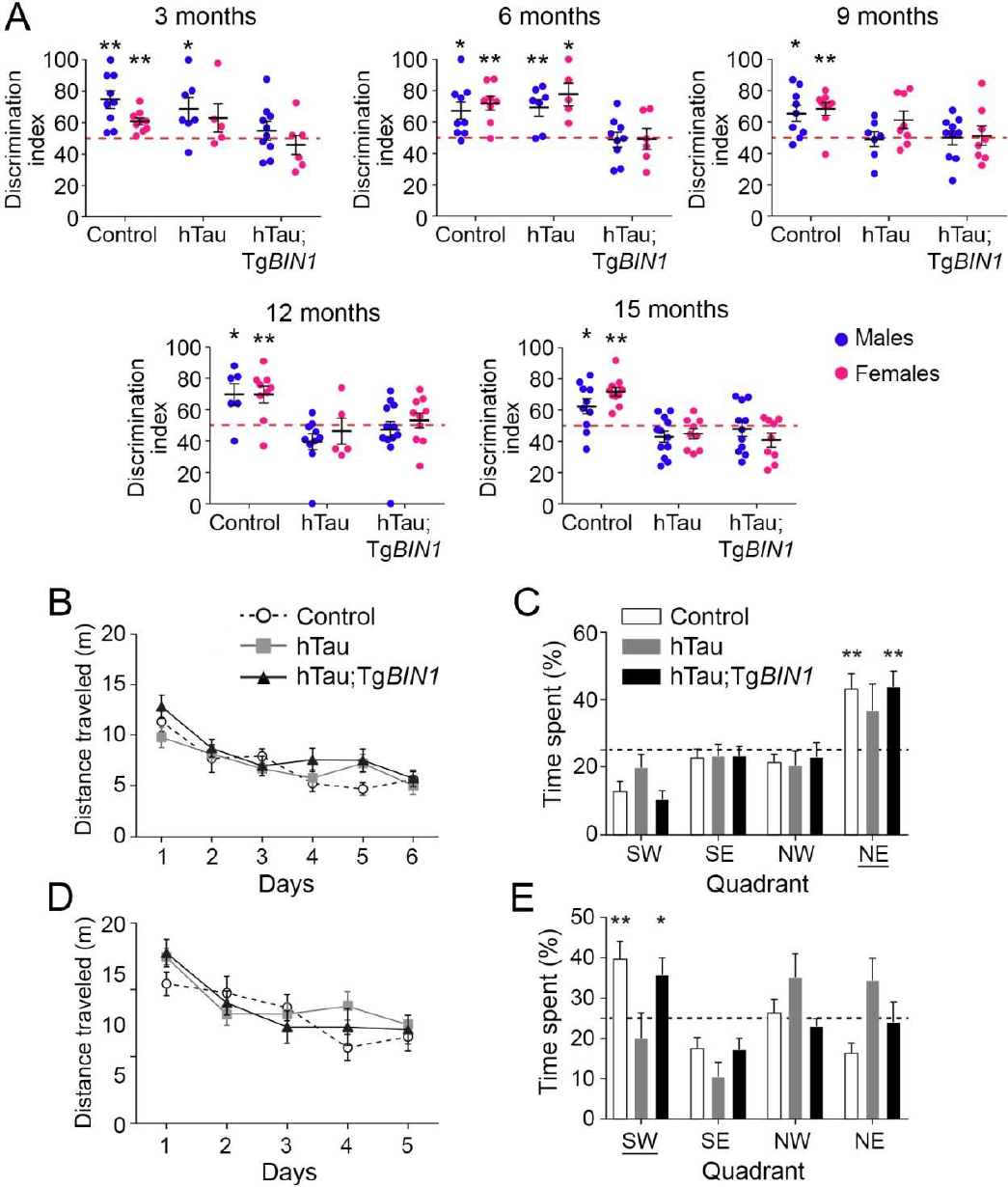
*BIN1* overexpression worsens hTau phenotypes in short-term memory and rescues long-term memory deficit due to MAPT overexpression in hTau males. **A.** Discrimination indices for novel object recognition with one hour of retention at 3, 6, 9, 12, and 15 months are shown for control, hTau, and hTau;Tg*BIN1* mice. Dashed lines represent object preference by chance. Blue dots, males; pink dots, females. One-sample t-test compared to chance at 50% *p<0.05, **p<0.01. **B.** Distance traveled to reach the platform of the Morris water maze for 12-month-old hTau and hTau;Tg*BIN1* males. Data represent mean ± SEM for consecutive days of acquisition (control, n=11; hTau, n=11; hTau;Tg*BIN1*, n=13). **C.** Probe test without platform at 12 months, performed 24 h after the last training session. Dashed line represents chance. Data represent mean ± SEM for each quadrant (control, n=11; hTau, n=11; hTau,Tg*BIN1*, n=13). Underlined quadrant marks original platform location. **D.** Distance traveled to reach the platform for 15-month-old hTau and hTau;Tg*BIN1* males. Data represent mean ± SEM for consecutive days of acquisition (control, n=11; hTau, n=10; hTau;Tg*BIN1*, n=13). **E.** Probe test without platform at 15 months, performed 24 h after the last training session. Dashed line represents chance. Data represent mean ± SEM for each quadrant (control, n=11; hTau, n=10; hTau,Tg*BIN1*, n=13). Underlined quadrant marks original platform location. One-sample t-test compared to chance at 25%; *p<0.05, **p<0.01.

In parallel to the NOR test, we assessed in this mouse cohort (non-naïve animals) the effect of *BIN1* and *MAPT* overexpression on long-term spatial memory using Morris water maze (MWM) tasks at the same relative ages. All groups were able to achieve the same performance in reducing the distance needed to reach the hidden platform (Fig. 1B-E and S3). The hTau mice displayed a deficit in recalling the platform location 24 h after the last training session by 12 months (Fig. 1B-E and S4). However, hTau;Tg*BIN1* males were able to perform this task at all ages tested up to 15 months, indicating that *BIN1* overexpression rescued the long-term and spatial memory of the hTau mice (Fig. 1B-E). The hTau;Tg*BIN1* females displayed a delayed deficit at 15 months compared to the hTau mice (Fig. S4). Notably, 15-month-old Tg*BIN1* mice did not have a deficit in this task (Fig. S5). To validate that the memory deficit observed for hTau mice were not due to a visual or locomotor deficit, we measured the distance and time required by the 15 month old mice to reach the visible platform. No difference was noted in the swimming velocities of different genotypes (Fig. S6). Overall, *BIN1* overexpression modulates hTau phenotypes by exacerbating short-term memory deficits and preventing long-term memory deficits.

### Human BIN1 expression prevents Tau intracellular inclusions and increases BIN1-Tau complexes in the hippocampus

The hTau mice have been described to develop detectable Tau aggregation and intracellular inclusions in the hippocampus and entorhinal cortex by 9 months.^27, 29^ We therefore tested the hypothesis that the mechanism underlying the rescue of the long-term and spatial memory deficits in hTau males through *BIN1* overexpression may be linked to an alteration of this aggregation. We sacrificed our cohort at 18 months and performed immunolabeling with antibodies specifically targeting Tau phosphorylation at both Ser202 and Thr205 (AT8 antibody) and at Thr231 (AT180 antibody) in the hippocampus (Fig. 2). As expected, no staining was evident in control mice. In hTau mice, Tau was mislocalized to the somatic compartment and formed prominent intracellular inclusions in the hippocampus (dentate gyrus, CA3, CA2, and CA1) (Fig. 2A). However, in hTau;Tg*BIN1* mice the number of cells with intracellular inclusions decreased by 5.9-fold or by 4.3-fold in the hippocampus when labeled with AT8 or AT180 antibodies, respectively (Fig. 2A-C). Since it is known that hyperphosphorylation of soluble Tau precedes Tau somatic inclusion,^30^ we determined if reduction of Tau inclusions upon BIN1 overexpression is due to an alteration of Tau phosphorylation pattern or of soluble Tau levels. However, no difference in soluble phosphorylated Tau protein was observed between hTau and hTau;Tg*BIN1* mice in the hippocampus (Fig. S7), indicating that BIN1 does not potentially regulate the level of soluble phosphorylated Tau protein or its phosphorylation pattern.

**Figure 2.**
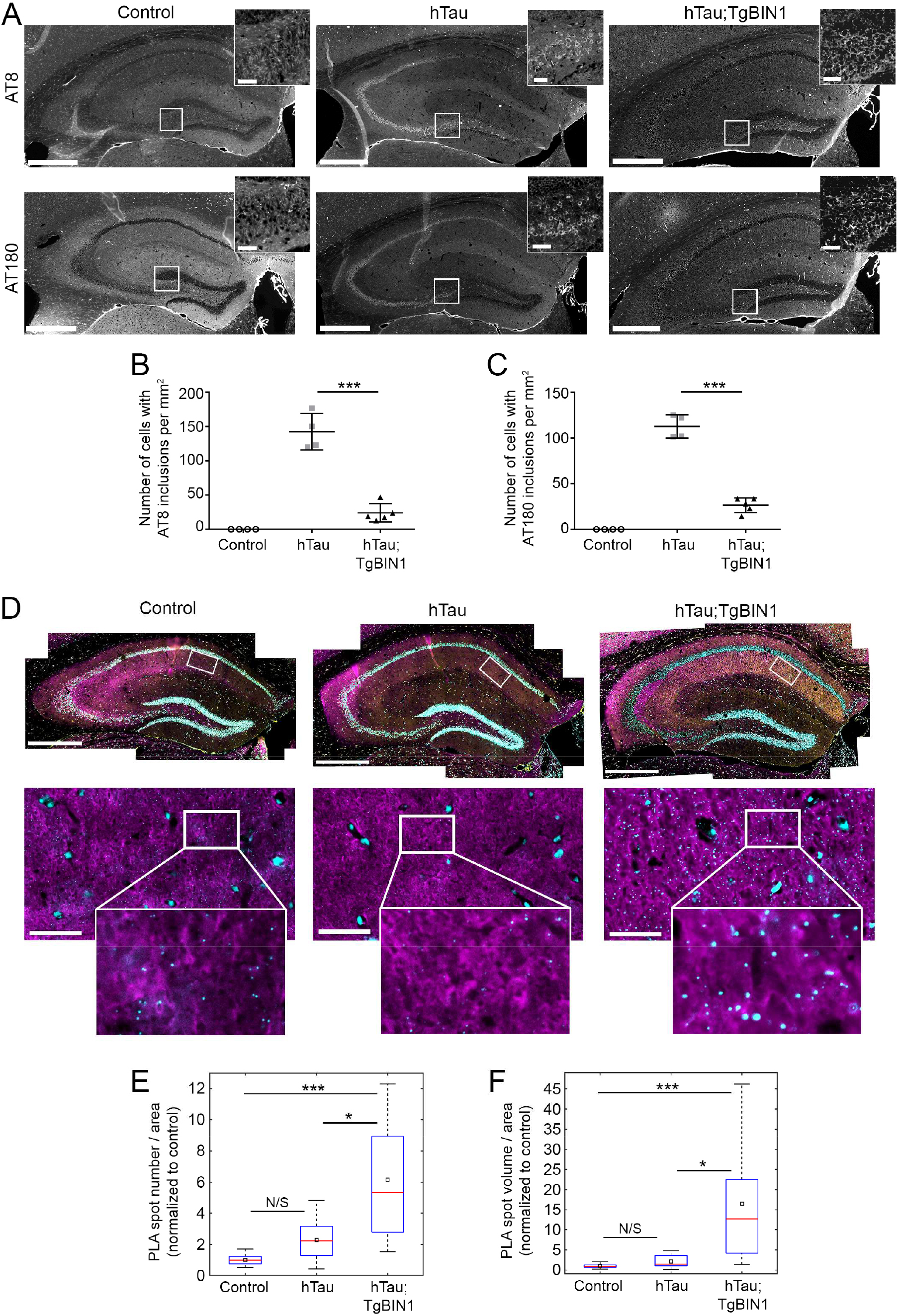
BIN1 overexpression prevents Tau inclusions and increases BIN1-Tau interaction in hTau hippocampi. **A.** Immunohistofluorescence of different phospho-Tau proteins in hippocampi of control, hTau and hTau;Tg*BIN1* males at 18 months. Antibodies used were detecting p-Ser202/p-Thr205 Tau (AT8) or p-Thr231 Tau (AT180). Insets show zooms of the hilus areas encompassing the neuronal cell bodies; intracellular inclusions are visible for hTau, but barely for hTau;Tg*BIN1*. Scale bars = 500 μm; insets, 50 μm. **B-C.** Quantification of the number of cells with intracellular Tau inclusions per mm^2^ in control, hTau and hTau;Tg*BIN1* mice labeled with the two phospho-Tau antibodies (control, n=4; hTau, n=4; hTau;Tg*BIN1*, n=5). **D.** BIN1-Tau PLA (cyan), and BIN1 (yellow), Tau (magenta), and Hoechst (white) stainings in the hippocampi of the same mice. Zoomed areas show PLA and Tau channels only. See Fig. S8 for Tubulin-Tau PLA, conducted as technical control. **E-F.** Quantification of BIN1-Tau PLA density. Data expressed as PLA spot number per tissue area (E) or total PLA spot volume per tissue area (F), normalized with control mean (control, n=9; hTau, n=11; hTau;Tg*BIN1*, n=12 hemispheres for spot number; control, n=10; hTau, n=12; hTau;Tg*BIN1*, n=12 hemispheres for volume). Red bars and black squares indicate sample median and mean, respectively. Kruskal-Wallis ANOVA, followed by multiple comparisons test with Tukey-Kramer correction; *** p < 0.0001; * p < 0.05. N/S, not significant. Scale bars = 500 μm; zooms, 50 μm.

It has been previously described that BIN1 is able to physically interact with Tau.^17, 31^ We assessed if BIN1 overexpression altered the amount and/or localization of BIN1-Tau complexes. For this purpose we used proximity ligation assay (PLA) in brain slices from sacrificed animals (Fig. 2D) and quantified the PLA density as a read-out of the BIN1-Tau interaction. We observed a strong increase in the PLA signal for the hTau;Tg*BIN1* mice when compared to both hTau mice and controls (2.7-fold and 6.2-fold in spot density, respectively) (Fig. 2D-F). As a positive control, we also used PLA to assess the interaction between α-tubulin and Tau and detected an increase in this interaction in hTau and hTau;Tg*BIN1* mice relative to controls (Fig. S8). Taken together, these data indicate that BIN1 overexpression increases the amount of BIN1-Tau complexes in the hippocampus and prevents Tau mislocalization and somatic inclusion, notably in the brain regions involved in long-term and spatial memory.

### BIN1 expression in neurons modulates BIN1-Tau interaction

Our data in transgenic mice support the idea that the BIN1-Tau interaction is relevant for the pathophysiological functions of Tau in AD and potentially in neurons. To gain further insight into the regulation of BIN1-Tau interaction, we monitored its dynamics during neuronal maturation in hippocampal primary neuronal cultures (PNC) at 7, 14, and 21 days *in vitro* (DIV), using western blots and PLA (Fig. 3). We first observed an increase in BIN1 and Tau amounts with time (Fig. 3A-B), whereas Tau phosphorylation was lower at certain epitopes, in particular, at Thr231 (Fig. 3C). Of note, this phosphorylation site has been described to inhibit the interaction between Tau’s proline-rich domain (PRD) and BIN1’s SH3 domains.^31^ The relative density of BIN1-Tau PLA volumes in the neuronal network was highly variable at DIV7 due to the low network density and it decreased with neuronal maturation (Fig. 3D-E), and the BIN1-Tau PLA signal was highly correlated with Tau volume irrespective of DIV (Fig. 3F), suggesting a uniform distribution of PLA signals in the network. We then assessed the impact of BIN1 expression on the PLA signal at DIV14, by downregulating BIN1 or overexpressing BIN1 neuronal isoform 1 (BIN1iso1) at DIV8 *via* transduction of lentiviruses expressing shRNA against BIN1 or the corresponding cDNA, respectively (Fig. 3G-I and S9). BIN1 downregulation led to a decrease in PLA signal; conversely, BIN1iso1 overexpression led to an increase in PLA signal (Fig. 3G and 3I). These data indicate that even if the BIN1-Tau interaction occurred at restricted loci in neurons (*e.g.*, at microtubule tips, as previously described ^31^), the BIN1-Tau complex formation depends on the global amount of BIN1 in neurons, as observed in the transgenic mice. Together, our data support the notion that variation in BIN1 expression affects the dynamics of BIN1-Tau complexes and their subsequent physiological and/or pathophysiological functions.

**Figure 3.**
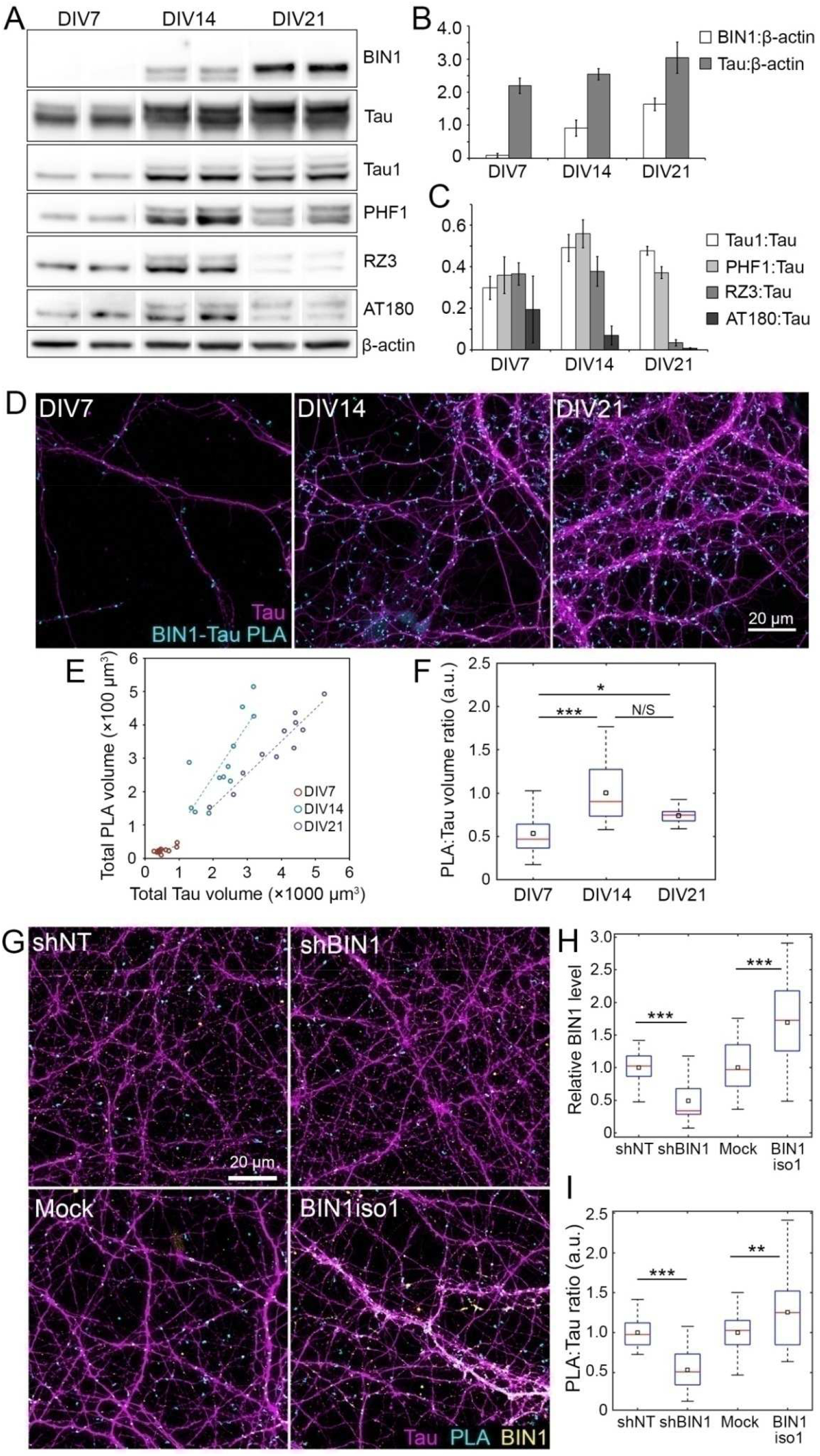
Characterization of BIN1-Tau interaction in primary neuron cultures (PNC). **A.** Representative immunoblots from neuronal extracts obtained at DIV7, DIV14, and DIV21 (in duplicate) showing BIN1 and total and phosphorylated forms of Tau (Tau1 for non-phospho Ser195/Ser198/Ser199/Ser202; PHF1 for p-Ser396/Ser404; RZ3 and AT180 for p-Thr231). **B-C.** Relative changes in BIN1 and Tau protein levels and in Tau phosphorylation during neuronal maturation. **D.** Representative images of PNC showing PLA spots and Tau immunolabeling during neuronal maturation. **E.** Change in PLA density during neuronal maturation. N = 3 independent experiments. **F.** Correlation between total PLA volume and total Tau volume obtained from a representative experiment. Each dot represents a confocal image. **G.** Representative images of PNC under- and overexpressing BIN1, showing PLA and Tau and BIN1 immunolabeling. shNT: non-targeting shRNA. **H-I.** Total BIN1 volume and PLA density in PNC under- and overexpressing BIN1, normalized with respective controls (shBIN1 with shNT and BIN1iso1 with Mock). N = 3 independent experiments. In box plots, red bars and black squares indicate sample median and mean, respectively. Wilcoxon rank-sum test; * p < 0.05; ** p < 0.01; *** p < 0.001; N/S: not significant.

### Identification of signaling pathways modulating the BIN1-Tau interaction in neurons

In addition to the BIN1 expression level as a modulator of the BIN1-Tau interaction, we had previously shown that phosphorylation of the Tau PRD domain (mainly at T231) inhibits its interaction with the BIN1 SH3 domain.^31^ This suggested that BIN1-Tau interaction dynamics likely depends on specific signaling pathways that regulate Tau phosphorylation. However, the cell signaling pathways susceptible to modulate the dynamic BIN1-Tau interaction remained unknown. To answer this question, we developed an agnostic strategy and set-up a semi-automated high-content screening (HCS) approach, using PNC as cellular model and PLA volume as readout for BIN1-Tau interaction (Fig. 4A).

**Figure 4.**
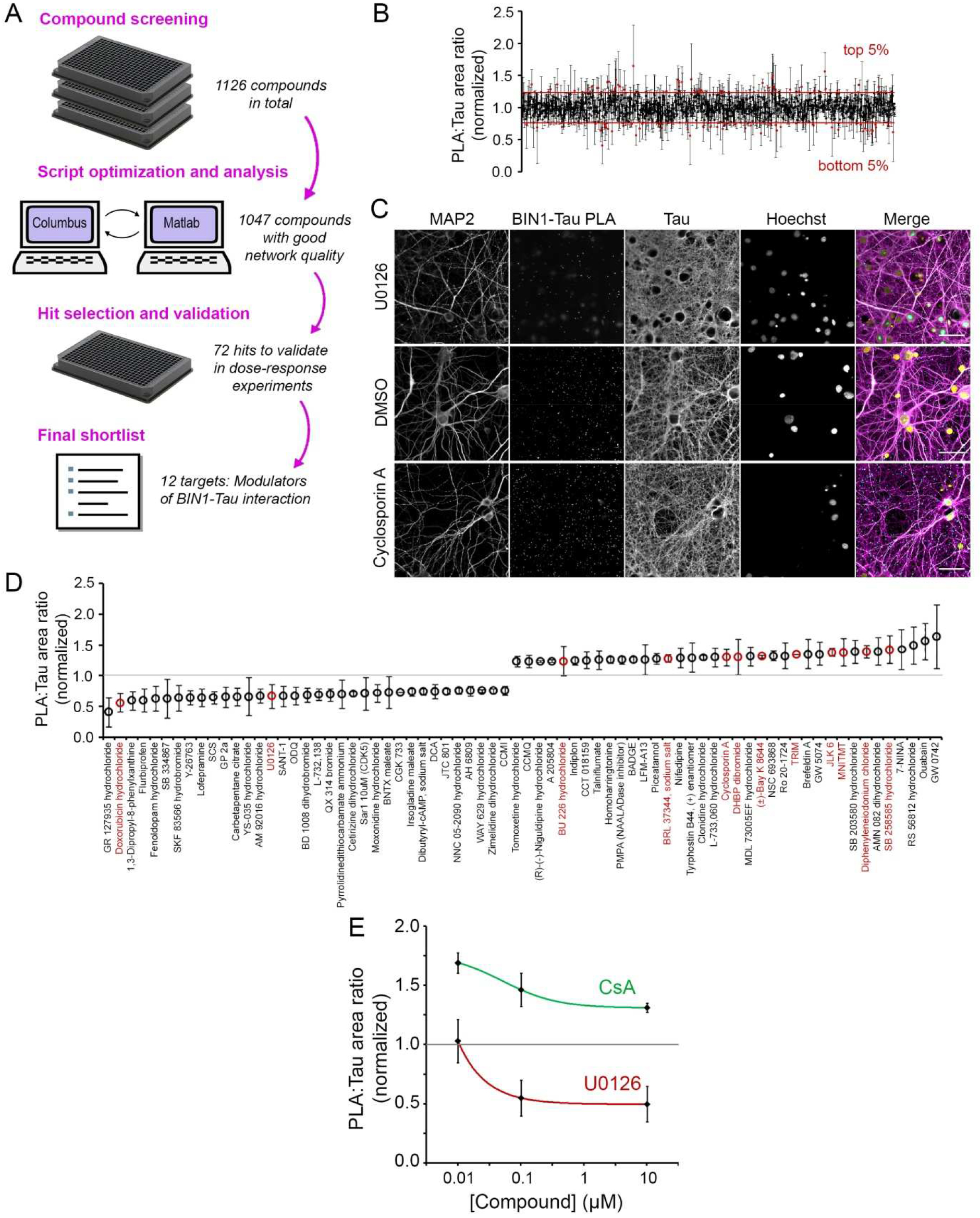
High-content screening (HCS) with PLA:Tau volume ratio in the Tau network as readout identifies the regulators of the BIN1-Tau interaction. **A.** The HCS workflow consists of compound screen (DIV21; 10 μM; 2.5 h) in PNC cultured in 384-well plates, plate-by-plate image segmentation and analysis, hit selection, and hit validation *via* dose-response experiments. **B.** Exemplary images from the HCS showing U0126 and Cyclosporin A (CsA) that decreased and increased PLA density, respectively. Scale bars = 50 μm. **C.** PLA:Tau area ratio for 1,047 compounds that did not induce damage in the neuronal network. Mean ± SD from 3 independent screens. **D.** Top and bottom 5% modulators (72 compounds) were retained for dose-response experiments. 12 compounds were validated in dose-response experiments are shown in red. **E.** Dose-response curves of U0126 and CsA (see Fig. S10 for all validated compounds). Mean ± SD from 3 independent experiments.

We tested a library of 1,126 compounds (at 10 μM) known to mainly target key elements of canonical pathways (see the Materials and Methods section for a full description of the HCS design). In brief, HCS was made in triplicate (one well per compound in each screen) using independent cultures. 79 compounds showed potential toxicity, as assessed by Tau and MAP2 network densities (Fig. 4B), and were excluded. We then applied several selection criteria to identify most promising compounds: (i) only compounds showing an effect in the same direction in all three independent screens were retained for further investigation; (ii) we selected the 10% of compounds showing the strongest variations (5% increasing PLA and 5% decreasing PLA). This led to 72 compounds for validation in dose-response experiments (Fig. 4C). Following this validation step, we were able to retain 12 compounds (marked red in Fig. 4D) that consistently exhibited the strongest variations in PLA signals. We grouped the targets of these compounds into 5 categories: (i) phosphorylation; (ii) nitric oxide synthase; (iii) Ca^2+^ homeostasis; (iv) membrane receptors; and (v) others (see Fig. S10 for the dose-response curves). As BIN1-Tau interaction has been shown to be modulated by phosphorylation,^31^ we decided to focus on two compounds whose targets are regulators of phosphorylation: (i) the Calcineurin (CaN) inhibitor Cyclosporin A (CsA), which, at 10 nM, increased PLA:Tau ratio by 42.6%; and (ii) the MEK inhibitor U0126, which, at 10 μM, decreased PLA:Tau ratio by 36.2% (Fig. 4E). In conclusion, our results show that CaN and MEK-dependent signaling pathways – among others – are able to modulate the complex dynamics of the BIN1-Tau interaction in neurons.

### The conformational change in BIN1 neuronal isoform 1 upon phosphorylation modulates BIN1-Tau interaction

Of particular interest, CaN is a Ser/Thr phosphatase which has been described to dephosphorylate Amphiphysin 1 (AMPH1), the homolog of BIN1.^32^ We thus postulated that CaN may also target BIN1 and sought potential phosphorylation sites within BIN1 explaining the increase in the PLA signal observed after CaN inhibition. Interestingly, we had previously characterized a conformational change in BIN1iso1 between open and closed forms. This involves an intramolecular interaction between the SH3 and CLAP PRD of BIN1iso1, making the SH3 domain unavailable for intermolecular interactions for instance with Tau.^26^ Since phosphorylations in the PRD have already been described to inhibit PRD/SH3 domains,^31^ we postulated that phosphorylation in the CLAP PRD domains of BIN1iso1 may favor BIN1’s open form and increase the BIN1-Tau interaction and consequently the PLA signal. When the protein sequences of AMPH1 and BIN1 are compared, their CLAP PRD domains appear to be highly conserved (Fig. 5A). Considering that AMPH1 T310 (corresponding to BIN1 T348) has been described to be phosphorylated by Cdks,^33^ we hypothesized that T348 (in the vicinity of the PRD sequence interacting with the BIN1-SH3 domain ^26^) may be controlling the open/closed conformation of BIN1iso1.

**Figure 5.**
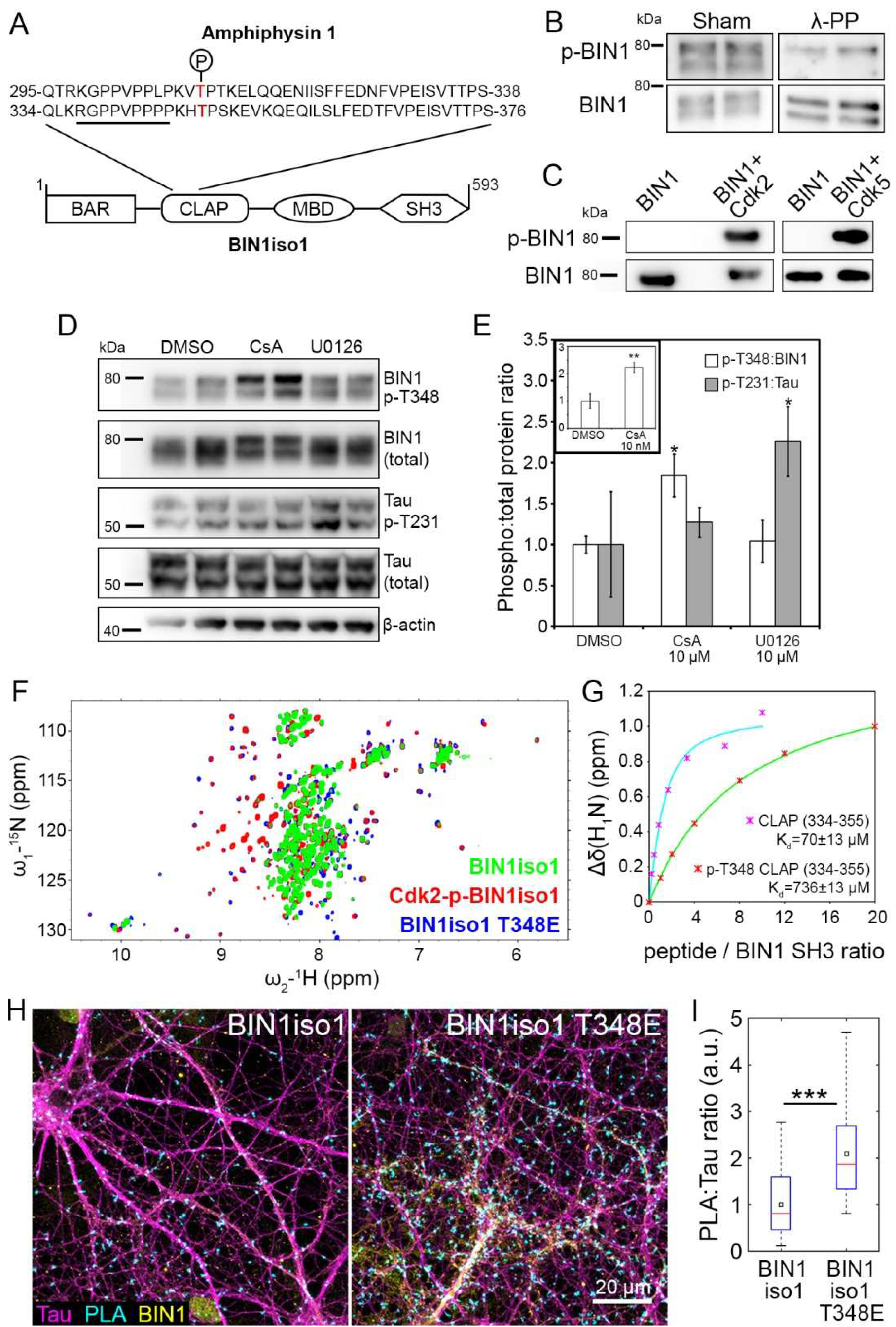
BIN1 phosphorylation at T348 regulates BIN1-Tau interaction by modulating open/closed conformation of BIN1. **A.** Alignment of Amphiphysin 1 and BIN1iso1; domains not to scale. The underlined sequence indicates the BIN1 PRD sequence interacting with the BIN1 SH3 domain. **B.** Lambda protein phosphatase (λ-PP) treatment dephosphorylates BIN1; 2 lanes per condition. **C.** *In vitro* phosphorylation assays with recombinant proteins show that Cdk2 and Cdk5 phosphorylate BIN1 at T348. Also see Fig. S12. **D-E.** Immunoblots and quantification showing the effects of U0126 and CsA (10 μM; 2.5 h) on BIN1 and Tau phosphorylation. Inset shows the effect of 10 nM CsA on BIN1 phosphorylation. Mean ± SD from 3 independent experiments. One-way ANOVA and paired t-test; * p < 0.05; ** p < 0.01. **F.** Behavior of BIN1-SH3 domain in the whole BIN1 isoform 1 protein as a function of phosphorylation by Cdk2 or of a mutation at threonine (T) 348 to glutamate (E) as monitored by ^1^H-^15^N HSQC spectra of BIN1iso1 CLAP T348E protein (in blue), Cdk2-phospho-BIN1iso1 (superimposed in red), and BIN1iso1 protein (superimposed in green). Also see Fig. S13. **G.** Titration of BIN1-SH3 domain with concentration of CLAP (334-355) or phospho-T348 CLAP (334-355) peptides. Normalized saturation curves (shown for residue 559), built from the gradual chemical shift changes (normalized; 1 denotes the largest change), are shown as pink stars for CLAP (334-355) and red stars for phospho-CLAP (334-355). Saturation curves are in cyan and green for CLAP (334-355) and phospho-CLAP (334-355), respectively. Also see Fig. S14. **H.** Representative images of PNC overexpressing BIN1iso1 and the BIN1iso1 T348E, its systematically open form, showing PLA signals and Tau and BIN1 immunolabeling. **I.** PLA density after normalization with respective BIN1 immunofluorescence in PNC overexpressing BIN1iso1 and BIN1iso1 T348E (for clarity, datasets were further normalized with the mean of BIN1iso1). N = 3 independent experiments. Red bars and black squares indicate sample median and mean, respectively. Wilcoxon rank-sum test; *** p < 0.001.

We first developed an antibody against BIN1 phosphorylated at T348 to determine if the BIN1 T348 phosphorylation occurred in neurons. Treating neuronal protein extracts with a protein phosphatase pool decreased BIN1 T348 phosphorylation (Fig. 5B). As control, Tau T231 phosphorylation was also decreased (Fig. S11A). Next, since T348 is within a consensus sequence recognized for phosphorylation by cyclin-dependent kinases (Cdks), we tested if Cdks were able to phosphorylate BIN1 T348. By using recombinant Cdk2 or Cdk5 and BIN1iso1, we showed that both kinases are able to directly phosphorylate T348 (Fig. 5C), as well as Tau T231 *in vitro* (Fig. S11B) confirming previous results^31^. We finally tested CsA and U0126 in PNC for their effect on BIN1 T348 and Tau phosphorylation. We observed that CsA – but not U0126 – was able to significantly increase BIN1 T348 phosphorylation in PNC (85±26% vs. 4±26%, respectively) suggesting that CaN is indeed able to dephosphorylate BIN1 at T348 (Fig. 5D-E). Remarkably, CaN inhibition did not impact Tau T231 phosphorylation, which we had previously described as a major modulator of the BIN1-Tau interaction,^31, 34^ suggesting that the BIN1 T348 phosphorylation alone drives the impact of CsA on PLA. Conversely, U0126 likely modifies the BIN1-Tau interaction through Tau T231 phosphorylation, without any impact on BIN1 T348 (Fig. 5D-E). Notably, we had previously characterized T231 as one of the 15 Ser/Thr sites where Tau gets phosphorylated by ERK2, downstream of MEK.^35^

To determine if phospho-T348 may control the dynamics of the open/closed conformation of BIN1iso1, we used nuclear magnetic resonance (NMR). We first tested whether this phosphorylation could impact the intramolecular interactions of BIN1 SH3 in the context of full BIN1iso1 protein. Signal from the BIN1-SH3 domain was observed in the spectra of Cdk2-phosphorylated recombinant BIN1iso1, whereas these same signals were barely detectable in the spectra of non-phosphorylated BIN1iso1 under identical acquisition and processing conditions (Fig. S12). Detection of these signals in the context of the large BIN1iso1 protein showed that the BIN1-SH3 domain kept some mobility and that the equilibrium was less in favor of the intramolecular interaction once the BIN1-CLAP domain was phosphorylated compared to the non-phosphorylated BIN1iso1 protein. However, since we detected multiple phosphorylation sites in the Cdk2-BIN1iso1 by NMR (Fig. S12), we generated a recombinant BIN1iso1 with T348E (BIN1-CLAP-T348E) to mimic the single phosphorylation event. Signals from the BIN1-SH3 domain were also detected in the spectra of the mutated BIN1iso1 T348E (Fig. S13), suggesting that phosphorylation at T348 is sufficient to shift to the BIN1iso1 open form (Fig. 5F). Finally, to further validate this observation, ^15^N-labeled BIN1 SH3 domain was titrated with CLAP (334-355) or phospho-CLAP (334-355) peptides and the titration was monitored using ^1^H-^15^N heteronuclear single quantum coherence (HSQC) spectroscopy of ^15^N-BIN1 SH3, one spectrum being recorded at each titration point (Fig. S14). The K_d_ values, obtained by fitting the chemical shift values measured in the spectra series to the saturation equation, were 71±13 μM for CLAP (334-355) peptide and 736±70 μM for phospho-CLAP (334-355) peptide, showing a 10-fold increase in K_d_ due to a single phosphorylation event in the peptide (Fig. 5G). Cumulatively, these results indicate that phosphorylation of T348 in the BIN1 CLAP domain is able to shift the dynamic equilibrium of the BIN1iso1 conformation towards the open form, thereby increasing the availability of the BIN1 SH3 domain for other interactions.

We next assessed whether the open/closed dynamics may impact the formation of the BIN1-Tau complex by controlling the availability of the BIN1iso1 SH3 domain in neurons and thus its ability to interact with Tau. For this purpose, we transduced at DIV8 hippocampal PNC with lentiviruses overexpressing wild-type BIN1iso1 and its mutated form, BIN1iso1-T348E, which, as previously demonstrated, leads to a systematically open form of BIN1iso1. We observed a 2.1-fold increase in PLA volume in PNC transduced with BIN1iso1-T348E when compared to BIN1iso1 (after normalization with respective BIN1 immunofluorescence) (Fig. 5H-I). This observation is in accordance with the increased availability of the BIN1iso1-T348E SH3 domain for Tau.

Finally, we quantified the amount of total and phospho-BIN1 (T348) neuronal isoforms of in protein extracts from 14 brain samples with increasing neurofibrillary pathology (Braak stages 0 to 6). The relative amounts of total and phosphorylated BIN1 exhibited a trend to decrease with increasing Braak stage (Fig. 6A-C). Surprisingly, the phospho-Bin1:BIN1 ratio exhibited a trend to increase with increasing Braak stage (Fig. 6D). Among the 14 individuals, 4 were controls and 12 were diagnosed with AD. After stratification based on the AD status, we observed a statistically non-significant decrease in total BIN1 in AD cases compared to controls (p = 0.05), but not in phospho-BIN1 (p = 0.71) (Fig. 6E-F). Interestingly, phospho-BIN1:BIN1 ratio was significantly increased in the brains of AD cases (p = 0.02) (Fig. 6G). Altogether, these data indicate that, in pathological conditions, the global level of the neuronal isoform of BIN1 is decreased, but a higher fraction of this BIN1 population is phosphorylated.

**Figure 6.**
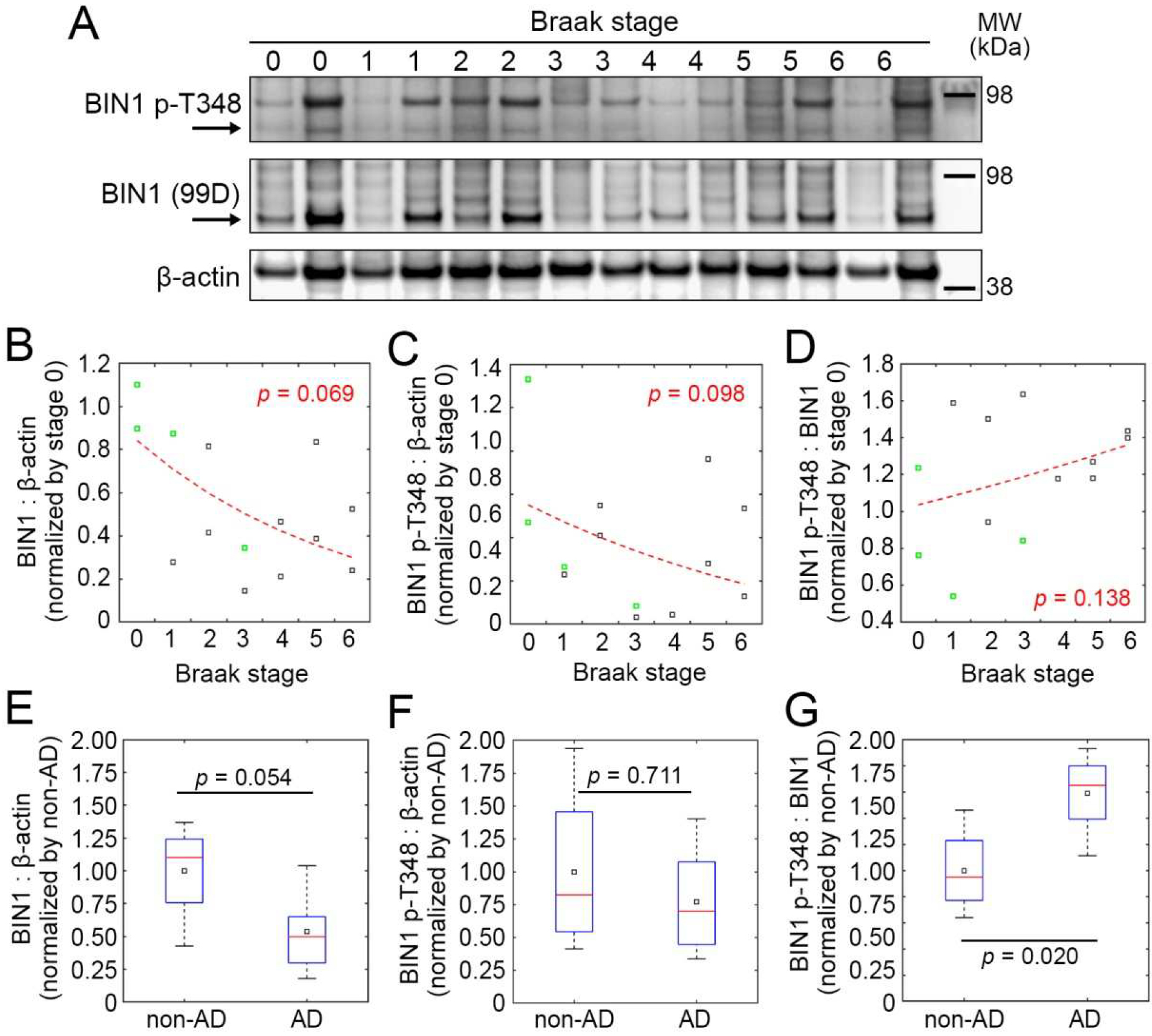
BIN1 amount and phosphorylation status in post-mortem AD brains. **A.** Western blots showing total BIN1 (99D antibody), BIN1 phosphorylated at T348 (p-T348), and β-actin in the temporal lobes of 14 individuals with increasing neurofibrillary pathology (Braak stage; see Table S3 for demographic details and pathological statuses). **B-D.** Quantification of the BIN1:β-actin, BIN1-p-T348:β-actin, and BIN1-p-T348:BIN1 signals, normalized with the mean of the control group (Braak stage = 0). Dashed red lines indicate exponential fits; *p*-values refer to the Kolmogorov-Smirnov test for the normal distribution of residuals. Data marked in green indicate non-AD cases according to neuropathological diagnosis. **E-G.** Comparison of BIN1:β-actin, BIN1-p-T348:β-actin, and BIN1-p-T348:BIN1 signals between non-AD and AD cases. Red bars and black squares indicate sample median and mean, respectively; *p*-values refer to the Wilcoxon rank-sum test.

## Discussion

There is no longer any doubt that BIN1 is a major genetic risk factor for AD.^7^ However, as for other GWAS-defined genes, it is often difficult to determine the implication of such genes in pathophysiological processes (or even in physiological ones in organs of interest). In this study, we aimed to determine if the BIN1-Tau interaction is involved in the neuropathological process of a mouse tauopathy model and to decipher the cellular processes and signaling pathways potentially regulating it.

**Figure7.**
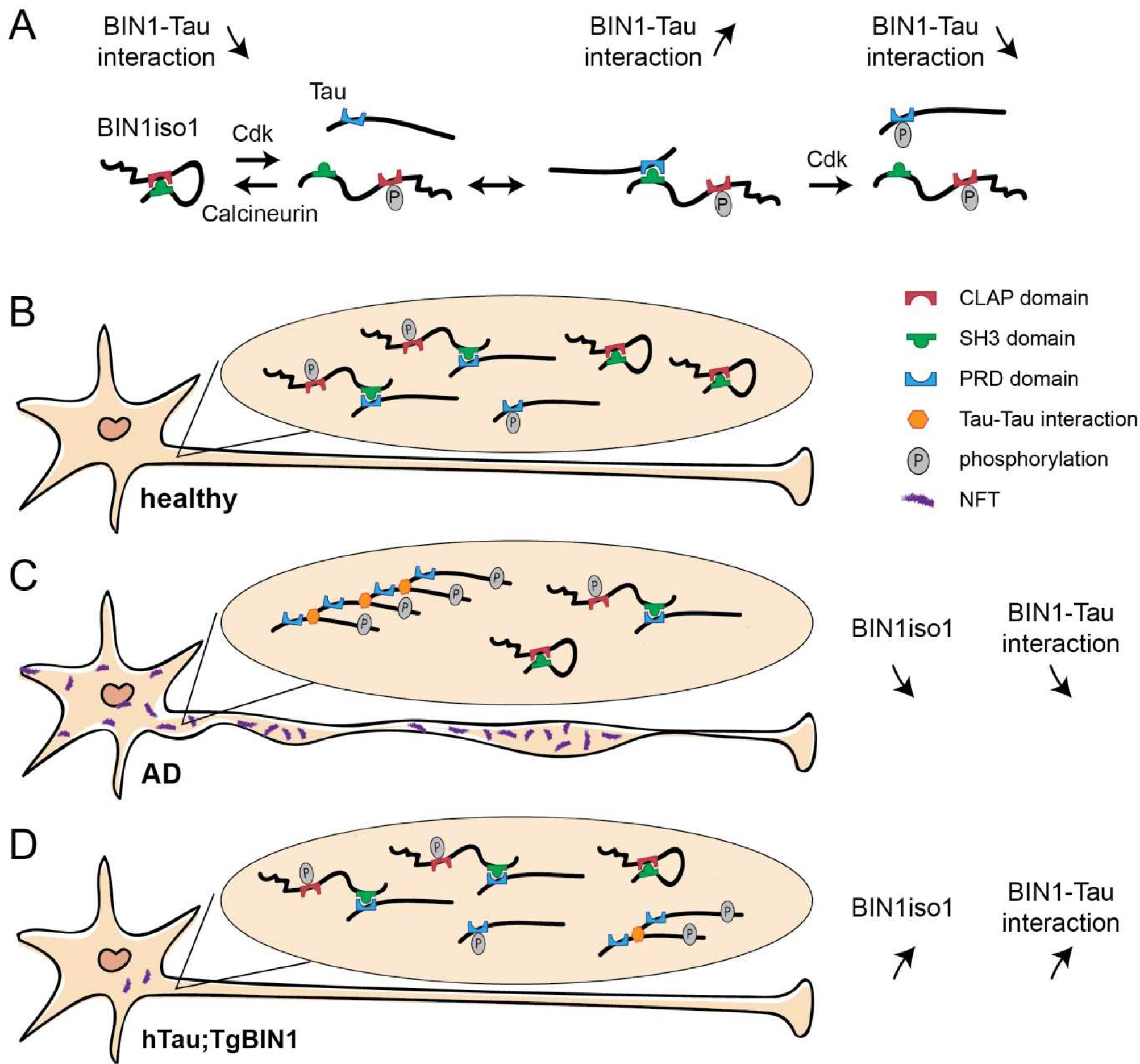
Complexity and dynamics of the BIN1-Tau interaction in neurons. **A.** At the molecular level, the open/closed conformation of BIN1 regulates the BIN1-Tau interaction in neurons under the control of the BIN1 T348 phosphorylation by CaN and Cdks. In addition, phosphorylation of Tau at T231 decreases the BIN1-Tau interaction. **B.** In healthy neurons, the BIN1-Tau interaction occurs at physiological levels. **C.** In AD pathology, a decrease in BIN1iso1 leads to a decrease in BIN1-Tau interaction, potentially favoring the formation of neurofibrillary tangles (NFT), in spite of potential compensation through BIN1 T348 phosphorylation. **D.** In tauopathy, overexpression of BIN1iso1 in neurons (as in the case for hTau;Tg*BIN1* mice) leads to an increase in the BIN1-Tau interaction and correlates with the disappearance of Tau somatic inclusions.

To determine if BIN1 could interfere with Tau pathology *in vivo*, we first developed a mammalian tauopathy model overexpressing BIN1 isoforms including neuron-specific forms in the brain. We observed that *BIN1* overexpression in the hTau mice expedited the appearance of short-term memory deficits from 9 to 3 months, but prevented spatial and long-term memory deficits up to 15 months, the highest age tested. Remarkably, the rescue of spatial and long-term memory by *BIN1* overexpression was associated with a strong increase in the BIN1-Tau interaction in the neuronal network and a strong decrease in phosphorylated Tau inclusions within the neuronal somata in the hippocampus. Next, we analyzed the BIN1-Tau interaction in the physiological context. BIN1 expression level appeared to be a strong modulator of the BIN1-Tau interaction in PNC. To identify signaling pathways modulating the BIN1-Tau interaction in neurons, we developed an agnostic HCS approach and determined a number of potential targets; one of best hits being an inhibitor of CaN, a Ser/Thr phosphatase. This observation led us to identify BIN1 phosphorylation at T348 as both a CaN target and a major regulator of the BIN1-Tau interaction. We determined that BIN1 phosphorylation at T348 increased the availability of the BIN1-SH3 domain to interact with Tau and consequently led to an increase in this interaction in neurons. Finally, we determined that neuronal BIN1 isoforms (mainly isoform 1) decreased in the brains of postmortem AD patients compared to control cases, whereas – surprisingly – phospho-BIN1(T348):BIN1 ratio increased, suggesting that this site may also be involved in the AD process. Overall we hypothesize that increased BIN1 expression and its phosphorylation on T348 protects hTau mice against spatial and long-term memory deficits (Fig. 7).

Altogether, our data support that a complex and dynamic regulation of the BIN1-Tau interaction is involved in the development of the AD pathophysiological process. However, the protective or deleterious effect of this interaction may vary depending on cognitive functions. Indeed, *BIN1* overexpression modulates *MAPT* phenotypes by exacerbating short-term memory deficits and by preventing long-term memory deficits. Both of these processes require the hippocampus, but the cortical regions involved are different, *i.e.*, lateral entorhinal cortex and medial entorhinal cortex, respectively.^36, 37^ The equilibrium between Tau and BIN1 levels may be slightly different in these cortical brain regions and in temporality, potentially explaining the opposite effects observed. In addition, signaling pathways controlling the phosphorylation of BIN1 and Tau, and subsequently the BIN1-Tau interaction may also differ temporally and regionally. However, since we developed a cohort study, it was not possible to evaluate such temporal and regional variations at each time of behavioral tests. It is nevertheless worth noting that the rescue of spatial and long-term memory by BIN1 overexpression was associated with a strong decrease in phosphorylated Tau inclusions within the neuronal somata and a strong increase in the BIN1-Tau interaction in the hippocampus. Remarkably, in hTau mice, the BIN1-Tau interaction was lower than in both control and htau;Tg*BIN1* mice. These observations thus suggest that the BIN1-Tau interaction may be protective by blocking the relocalization and accumulation of phosphorylated Tau in the neuronal somata, a major hallmark of AD.

The hypothesis that a dynamic regulation of the BIN1-Tau interaction is involved in AD process also implies that a high level of BIN1 expression would be protective. However, we previously found that total BIN1 mRNA is over-represented in the brains of AD cases compared to controls,^38^ but did not evaluate at that time the isoform-dependency of the BIN1 expression. Subsequent publications reporting protein levels showed that unlike the overexpression of ubiquitous isoforms, the neuronal isoforms were specifically underexpressed in the AD brains.^39, 40^ We validated this observation in brain samples and showed that this decrease was dependent on the Braak stage (Fig 6). Since the neuronal isoforms are the main isoforms that are overexpressed in the brain of our transgenic mice model,^21^ these data corroborate the idea that specific overexpression of the neuronal BIN1 isoforms may be protective. We may thus postulate that the overexpression of neuronal BIN1 isoforms in the Tg*BIN1* mouse reverses a neuropathological process that occurs in AD brains. This protective effect could be explained by the BIN1-Tau interaction in neurons. However, we cannot exclude other potential mechanisms. Indeed, we observed that at 18 months *MAPT* over-expression is associated with myelin abnormalities, and a significant rescue of this phenotype was observed in hTau;Tg*BIN1* mice (Fig. S15). Of note, the over-expression of BIN1 alone did not induce any myelin abnormalities (Fig. S16; also see supplementary results). Thus, the memory impairments observed in the behavioral analyses of the hTau mice may also be associated with myelin disorganization in the fornix, and be rescued upon BIN1 overexpression. Interesting, BIN1 has been described to be strongly expressed in oligodendrocytes ^22^ and Tau has been also previously linked with potential myelin dysfunction in tauopathies.^41^

Identifying the signaling mechanisms controlling the BIN1-Tau interaction is of high interest to understand the pathophysiological processes in AD. These pathways could be either protective or deleterious, by favoring or abrogating the BIN1-Tau interaction, respectively. In this report we characterized a key regulatory element, which is the phosphorylation of BIN1 at T348. Remarkably, we determined that the phospho-BIN1(T348):BIN1 ratio increased with increasing Braak stage in the brains of AD cases. These findings suggest that a higher fraction of brain BIN1 isoforms is phosphorylated at T348 in AD brains, where the global level of neuronal BIN1 isoforms is decreased. This may imply that the relative increase in BIN1 T348 phosphorylation occurs to compensate in part the decrease in the neuronal BIN1 isoforms in order to maintain the BIN1-Tau interaction. Altogether, these observations suggest that BIN1 T348 phosphorylation is involved in the development of AD.

Our data thus indicate that the BIN1-Tau interaction is complex and dynamic, potentially controlled by numerous actors modifying the level of phosphorylation of both BIN1 and Tau, including Cdks and CaN. Indeed, we had previously shown that the phosphorylation of Tau at T231 was a major regulator of the BIN1-Tau interaction, but in the opposite direction, *i.e.*, leading to a decrease in this interaction. Importantly, the increase in Tau phosphorylation at T231 is considered as an early marker of the development of AD.^30^ This dual BIN1/Tau regulation is illustrated in our HCS screening, which revealed that inhibiting CaN favors the BIN1-Tau interaction by increasing BIN T348 phosphorylation, whereas inhibiting MEK hinders it by increasing Tau T231 phosphorylation. Cdks – particularly Cdk5 – highlight this complexity, since these kinases are able to phosphorylate both BIN1 T348 and Tau T231, but with opposite effects on the abilities of Tau and BIN1 to interact with each other: increased Cdk5 activity would increase BIN1’s affinity for Tau through phosphorylation of BIN1 at T348, and, conversely, would decrease Tau’s affinity for BIN1 through phosphorylating Tau at T231 (Fig. 7). This complex interplay between actors modulating BIN1 and Tau phosphorylation may be a limitation for developing drugs to favor or prevent the BIN1-Tau interaction. A better understanding of the mechanisms involved will thus be needed to identify potential cell signaling pathways and drug targets that would uncouple the BIN1-Tau phosphorylation crosstalk. In this context, CaN-dependent pathways may be of therapeutic interest, since we observed that only BIN1 T348 is modulated by CaN, but not Tau T231.

In conclusion, we reveal the impact of overexpression of BIN1, a major genetic risk factor of AD, in a tauopathy model. Our data also reinforce the hypothesis that a potential protective impact of this overexpression on the AD process may be linked to the direct interaction of BIN1 and Tau, and depends strongly on the phosphorylation statuses of both proteins.

## Materials and Methods

### Animal ethics

Animal experiments were approved by the Com’Eth (project file: 2014-056) and accredited by the French Ministry for Superior Education and Research in accordance with the Directive of the European Parliament: 2010/63/EU. For all tests described, mice were kept in specific pathogen free conditions with free access to food and water, and were bred with littermates. The light cycle was controlled as 12 h light and 12 h dark (lights on at 7AM). Before all behavioral experiments, handling was done every day for one week before the beginning of the experiment.

### Mouse lines and genotyping

We used several mouse lines carrying the inactivation of *Mapt*: B6.Cg *Mapt^tm1(EGFP)Klt/+^*, noted here *Mapt^+/-^*,^42^ a line overexpressing human Tau: B6.Cg *Mapt^tm1(EGFP)Klt/tm1(EGFP)Klt^ Tg(MAPT)8cPdav/J*, named here hTau,^28^ and another line overexpressing human BIN1: B6 *Tg(Bin1)U154.16.16Yah*, named here Tg*BIN1*/0.^21^ In order to generate cohorts of animals carrying hTau alone, hTau;Tg*BIN1*, and *Mapt^+/-^* as control littermate, we crossed *Mapt^+/-^*;*Tg(MAPT)8cPdav/J* with *Mapt^+/-^*;Tg*BIN1*. All animals were crossed on C57BL/6J background. Primer sequences are available in Table S1.

### Design of behavioral experiments

Animals studied in behavioral tasks were both males and females. Same animals were longitudinally tested at 3, 6, 9, 12, and 15 months. All animals were killed at 18 months for histology and molecular biology experiments.

### Novel object recognition task

This task was performed in the same conditions as in the open field paradigm (see above). The objects to be discriminated were a glass marble (2.5 cm in diameter) and a plastic dice (2 cm). The animals were first habituated to the open field for 30 min. The next day, they were submitted to a 10 min acquisition trial during which they were placed in the open field in the presence of two similar objects (object A; marble or dice). The time the animal took to explore the object A (when the animal’s snout was directed towards the object at a distance ≤ 1 cm) was recorded manually. A 10 min retention trial was performed 1 h later. During this trial, one of the familiar objects in the open field was replaced with a new one (object B), and the time periods that the animal took to explore the two objects were recorded (t_A_ and t_B_ for objects A and B, respectively). Two exclusion criteria were applied to select those animals that had memorized the objects: (i) during the acquisition trial, mice exploration should be longer than 3 s, and (ii) during the retention trial, mice exploration should also be longer than 3 s. The exploration index for object B was defined as (t_B_ / (t_A_ + t_B_)) × 100. Memory was defined by the percentage of time animals spent investigating the novel object statistically different from the chance (50%). To control for odor cues, the open field arena and the objects were thoroughly cleaned with 50% ethanol, dried, and ventilated between sessions. All animals were tracked with Noldius software (Ethovision).

### Morris water maze task

The Morris water maze was used to test spatial learning and memory. Each session was performed one week after NOR task and constituted the last behavioral experiment. The water maze is a circular pool (150 cm in diameter, 60 cm in height), filled with water up to 40 cm mark that is maintained at 20-22°C, and made opa que using a white aqueous emulsion (Acusol OP 301 opacifier). The surface was split into 4 quadrants: South-East (SE), North-West (NW), North-East (NE), and South-West (SW). The escape platform, made of rough plastic, was submerged 1 cm below the water’s surface. Experiments were performed to study reference memory through a spatial search strategy that involved finding the hidden platform. The spatial memory session consisted of a 6-day (J1 to J6) learning phase with four 90 s trials per day. Each trial started with mice facing the interior wall of the pool and ended when they climbed onto the platform located on the SE quadrant, or after a maximum searching time of 90 s. The starting position was changed pseudo-randomly between trials. Mice were left undisturbed in their home cage for 90 min intertrial intervals. On the 7^th^ day, mice were given the 60 s probe test, in which the platform had been removed. The distances traveled in each quadrant (NW, NE, SW, and SE) were recorded, as well as the time spent in the target quadrant. At 6, 9 and 12 months of age, the platform was located in the NE quadrant, whereas at 15 months of age, the platform was located in the SW quadrant. All animals were tracked with Noldius software (Ethovision).

### Brain protein extraction and Western blotting

Mice were killed by cervical dislocation and brains were quickly removed and dissected. Structures were immediately frozen in liquid nitrogen, and conserved at -80°C. For protein extraction we used fresh extraction buffer with pH adjusted to 7,5 (20 mM Tris at pH = 7,5; 50 mM NaCl; 2 mM EGTA; 1% Triton X-100; 10 mM NaF; 1 mM Na_3_VO_4_; 2 mM β Glycerophosphate; cOmplete™ EDTA-free protease inhibitor cocktail). Tissues were lysed using Precellys apparatus and centrifuged at 33,000 ×g for 30 min. Protein quantification was performed using the BCA protein assay (Thermo Scientific; Waltham, MA). 10-20 μg of total protein from extracts were separated in SDS–polyacrylamide gels (10%) and transferred to nitrocellulose membranes. Depending on the target protein, we used bovine serum albumin or milk (5% in Tris-buffered saline with 0.1% Tween-20, TTBS; 1 h at RT) to block non-specific binding sites of phosphorylated and non-phosphorylated proteins, respectively. Immunoblotting was carried out with primary antibodies (Table S2) for 1 h at RT. Then membranes were washed 5 times in TTBS, followed by incubation with secondary antibodies conjugated with horseradish peroxidase (Table S2). Immunoreactivity was visualized using ECL chemiluminescence system (SuperSignal™, Thermo Scientific). Chemiluminescence was captured with Amersham Imager and signals were quantified with ImageJ (NIH; Bethesda, MD).

### Immunofluorescence in brain slices

Mice were anesthetized with 5% ketamine and 10% xylazine and perfused first with PBS and then with 4% paraformaldehyde (PFA) in PBS. After removal, brains were immerged in 4% PFA overnight at 4°C, followed by multiple rinses w ith PBS, and put in 30% sucrose in PBS until they sink. Once they sink, they were embedded in O.C.T. tissue freezing compound (Scigen; Gardena, CA), and stored at -80°C until th ey were cut with a cryostat at 10 μm thickness. For immunofluorosence, slices were first permeabilized with 0.1% Triton in PBS, with 10% horse serum and 5% BSA for 30 min. The primary antibody (Table S2) was then applied overnight at 4°C in the permeabilization bu ffer. After multiple rinses with PBS, the secondary antibody (Table S2) in 0.1% Triton was applied for 1 h at RT. After multiple rinses, slices were stained with 1:1000 Hoechst (Sigma; St. Louis, MO). After multiple rinses, slices were mounted in Fluorsave (Merck Millipore; Darmstadt, Germany). Slices were imaged with NanoZoomer slice scanner (Hamamatsu Photonics; Massy, France).

### Electron microscopy of brain slices

Mice were PFA-fixed as described. After removal, brains were immerged in 4% PFA and 4% glutaraldehyde in PBS overnight at 4°C. Coronal sec tions were obtained with Leica VT1000 vibratome (Leica Biosystems; Nanterre, France), and the tissue was cut to expose the dorsal fornix and the upper part of the hippocampus as shown in Fig. 2. The tissues were post-fixed in 1% osmium tetroxide, dehydrated through graded ethanol (50, 70, 90, and 100%) and propylene oxide for 30 min each, and embedded in Epon 812 (EMS; Hatfield, PA). Semithin sections were cut at 2 μm on an ultra-microtome (Ultracut UCT; Leica) and ultrathin sections were cut at 70 nm, contrasted with uranyl acetate and lead citrate, and examined at 70 kV using a Morgagni 268D electron microscope (Thermo Scientific). Images were captured digitally by Mega View III camera (Soft Imaging System; Münster, Germany).

### Primary neuronal culture

Culture media and supplements were from Thermo Scientific, unless mentioned otherwise. Primary hippocampal neurons were obtained from P0/P1 rats, according to previously described procedures ^43, 44^ with minor modifications. Briefly, cortices and hippocampi were isolated from new-born rats, washed with ice-cold dissection medium (HBSS supplemented with HEPES, sodium pyruvate, and penicillin/streptomycin), and trypsinized (2.5%; 10 min; 37°C). Trypsin was inactivated with dissociation me dium (MEM supplemented with inactivated FBS, Glutamax, D-glucose (Sigma), MEM vitamins, and penicillin/streptomycin), followed by DNase (5 mg/ml; Sigma) incubation for 1 min and wash with dissection medium. Media was replaced by dissociation medium and tissue was triturated with a fire-polished cotton-plugged Pasteur pipette to obtain a homogenous cell suspension, followed by centrifugation (200 ×*g* for 5 min) and wash with dissociation medium. Cells were resuspended in culture medium (Neurobasal A supplemented with Glutamax and B_27_ neural supplement with antioxidants), counted, and plated in 384-well plates (Greiner bio-one; Kremsmünster, Austria) at a density of 50,000 cells/cm^2^ for HCS, on Ø13 mm coverslips in 24-well plates at a density of 25,000 cells/cm^2^ for proximity ligation assay (PLA), or directly in 24-well plates without coverslips at density 100,000 cells/cm^2^ for immunoblots. Coverslips and plates were pre-coated with poly-L-lysine (Alamanda Polymers; Huntsville, AL) overnight at 37°C and rinsed thoroughly with water. After 20-24 h, culture media was replaced with supplemented Neurobasal A medium and cultures were maintained in a tissue culture incubator (Panasonic; Osaka, Japan) at 37°C and 5% CO_2_ for 7, 14, or 21 days.

### Viral transductions

PNC were transduced on DIV8 with lentiviral constructs for silencing (MOI = 4) using Mission pLKO,1-puro-CMV-shRNA vectors (Sigma), non-targeting (05191520MN) and shBIN1 (TRCN0000380439). Overexpression constructs were obtained from Gene Art (Thermo Fisher) based on pLenti6/Ubc/v5-DEST vectors (Life Technologies, Carlsbad, CA): BIN1iso1 (NM_009668), BIN1iso1 phosphomimetic T348E (cDNA with Thr^348^→Glu), BIN1 isoform 9 (NM_139349), and an overexpression control vector (mock). The transduction was performed according to a previously described procedure ^45^ with minor modifications: For PNC in 24-well plates, viral constructs at multiplicity of infection (MOI) 2 were added to pre-warmed supplemented Neurobasal A media with Polybrene (0.1% final concentration; Sigma) at 10× concentration. Half of the culture media from multi-well plates were collected and stored. The transduction mixture was added to each well to reach 250 μl final volume and neurons were incubated for 6 h. At the end of this period, wells were topped with 250 μl collected media and neurons were maintained in the incubator until fixation or protein harvest. Transduced neurons were either fixed or harvested on DIV14.

### Immunoblotting

PNC were harvested in minimum volume of 40 μl/well in ice-cold lysis buffer as described earlier.^38^ Lysates were mixed with 4× LDS (Novex; Life Technologies) and 10× reducing agent (Novex) loaded on pre-cast NuPage 4-12% bis-Tris acrylamide 10 well gels (Novex) and transferred to nitrocellulose membranes using the BioRad Trans-blot transfer system kit (BioRad, Hercules, CA). Membranes were blocked in 5% non-fat milk in 1x TNT buffer. Primary antibodies were diluted in SuperBlock T20 (TBS) blocking buffer (Thermo Fisher) and kept at 4°C overnight: mouse BIN1-99D (clone 99 D; 1:1,000; cat. no. 05-449, Merck Millipore), rabbit TauC (1:10,000), mouse beta-actin (1:10,000; Sigma), rabbit phospho-BIN1 Thr 348 (1:10,000; custom made by Biotem, Apprieu, France), mouse Tau 1 non-phospho Ser 195-Ser 202 (aa197-205) (1:10,000; Merck Millipore), mouse AT180 phospho Thr 231 (1:500, Thermo Fisher), mouse RZ3 Thr 231 (1:500), and mouse PHF1 phospho Ser396/404 (1:1000). The last two antibodies were kind gifts from Peter Davies. We further confirmed the specificity of this antibody for the neuronal isoform by silencing BIN1 and overexpressing BIN1iso1 or BIN1iso9 (Fig. S17). Detection was performed using horseradish peroxidase (HRP)-conjugated secondary antibodies (1:5000, Jackson) for 1-2 h at RT. The membrane was revealed through chemiluminescence (Luminata Crescendo^TM^, EMD Merck Millipore) and imaged with Amersham Imager 600 (GE Healthcare, Mississauga, Canada). The images were quantified with ImageQuantTL Software (GE Healthcare).

### Analysis of neuropathological human sample cohort

Assessment of AD-related neurofibrillary pathology (Braak stage) was performed for 14 individuals after death (Table S3) with immunostaining of paraffin sections with AT8 antibody, which detects hyperphosphorylated Tau.^46^ Protein extractions from the frozen temporal lobe tissue samples were performed as previously described.^47^ Protein quantification was performed using BCA protein assay. Total proteins (20 μg/lane) were separated on 4-12% Bis-Tris-polyacrylamide gel electrophoresis (PAGE; Invitrogen) under reducing conditions and subsequently blotted onto polyvinylidene difluoride membranes using iBlot 2 Dry Blotting System (Thermo Scientific). Primary antibodies against phospho-BIN1 Thr 348 (1:1,000), total BIN1 (1:1,000) and β-actin (1:1,000; cat. no. ab8226, Abcam) were used for immunoblotting. After incubation with the appropriate HRP-conjugated secondary antibodies, the protein bands were detected using ImageJ.

### Lambda protein phosphatase assay

Crude protein extracts were incubated with Lambda protein phosphatase (New England Biolabs; Ipswich, MA), following supplier’s instructions with minor changes. DIV21 PNC were harvested on ice in 40 μl ice-cold lysis buffer per well without protein phosphatase inhibitors, lysates were sonicated, centrifuged for 10 min at 1,000× *g* and the supernatant was distributed into 2 new tubes; volumes were adjusted to 40 μl with MilliQ H_2_O, and supplemented with 5 μl of 10× NEBuffer and 5 μl of 10 mM MnCl_2_ (provided with the enzyme); 1 μl of lambda protein phosphatase (λ-PP) was added to one of the tubes and both tubes were incubated for 30 min at 30°C. 4× LDS and 10× reducing agent were added to the tubes, samples were boiled at 95°C for 10 min and i mmunoblotted as described before.

### *In vitro* assay with recombinant proteins

BIN1 phosphorylation *in vitro* was assessed in kinase buffer containing 20 mM MOPS, pH 7.4, 5 mM MgCl_2_, 100 μM ATP, and 1 mM DTT. Purified GST-BIN1 (500 ng) was incubated with recombinant GST-tagged Cdk5/p35 (100 ng) at RT for 1h. The reaction was terminated by the addition of boiled SDS sample buffer. After electrophoresis of the samples were run on SDS-PAGE. In addition, Cdk2/CycA3 kinase ^48^ was used to obtain Bin1iso1 phosphorylated on T348 residue. The capacity of the kinase to phosphorylate T348 was first verified using the CLAP (334-355) peptide as substrate and mass spectrometry to assess the addition of a phosphate group. In addition, the phosphorylated peptide was detected using the antibody directed against pT348 (Fig. S12A, inset). For NMR experiments, 100 μM ^15^N-BIN1iso1 was incubated with recombinant Cdk2/CycA3 kinase (molar ratio 1/100), for 3 h at 37°C, in the presence of 2 mM ATP, 2.5 mM MgCl _2_, 2 mM EGTA, 2 mM DTT, 30 mM NaCl and protease inhibitors in 50 mM HEPES, pH 8.0 (Fig. S12). Control experiment was performed in the absence of ATP. Phosphorylation of Bin1Iso1 at T348 was verified using western blot analysis with an antibody directed against pT348.

### NMR spectroscopy

NMR experiments were recorded at 20°C on Bruker 900-MHz spectrometer. NMR measurements were performed in 50 mM sodium phosphate buffer, pH 7.3, 30 mM NaCl, 3 mM DTT and 10% D_2_O. BIN1iso1, BIN1iso1-CLAP-T348E and Cdk2-phospho-BIN1iso1 ^1^H-15N HSQC spectra were all recorded with a TXI probe at a protein concentration of 100 μM. These 2D spectra were acquired with 3072 points in the direct and 180 points in indirect dimensions for spectral width of 13 ppm and 26 ppm, respectively, and with 512 scans. BIN1-SH3 domain ^1^H-^15^N HSQC spectrum was recorded with a cryogenic probe with 3072 points in the direct and 256 points in indirect dimensions for spectral width of 14 ppm and 26 ppm, respectively, and with 48 scans. Spectra were processed using TopSpin software (Bruker). BIN1-SH3 domain backbone assignments were previously reported.^26^ The NMR titration data were obtained by adding aliquots of 4 mM stock solutions of unlabeled peptides Q L R K G P P V P P P P K H **T** P S K E V K Q CLAP (334-355) or phospho-T348 CLAP (334-355), phosphorylated residue in bold in the sequence, to 100 μM ^15^N-labeled BIN1-SH3 domain, using HSQC spectra to monitor changes in amide and tryptophan indole chemical shift values. K_d_ were calculated based on these data (see Supplementary Information for details).

### Semi-automated high-content screening for modulators of BIN1-Tau interaction

A compound screen was setup by combining a commercial library of 1,120 compounds (10 μM; #2890; Tocris Biosciences, Bristol, UK), 6 Sanofi proprietary compounds (0.1, 1, and 10 μM; Sanofi; Chilly-Mazarin, France), Okadaic acid (1 μM; Merck Millipore) as a control compound, and DMSO (0.1%; VWR; Radnor, PA). Tocriscreen™ Mini is a library of well-characterized biologically active compounds that allows the screening of a wide-range of cellular processes, such as inflammation, apoptosis, cell differentiation, signal transduction, intracellular transport. 1000× stock compounds were transferred into intermediate 384-well plates using Echo 550 liquid Handler (Labcyte; San Jose, CA), and plates were sealed and kept at -20°C. Neurons cultured in 384-well plates were maintained for 21 days and transferred to HCS platform incubator (Liconic instruments; Mauren, Liechtenstein) on the day of screening. Compounds in intermediate plates were resuspended in 30 μl Neurobasal A, to reach 5× concentration, followed by a 2 min-long centrifugation at 100 ×*g*. 10 μl of resuspended compounds were then added into respective wells in PNC plates using Bravo automated liquid handling platform (Agilent; Santa Clara, California, USA), containing 40 μl of culture media, and plates were returned to the incubator. To achieve equal treatment duration for all plates, the compounds were resuspended and transferred with 10 min intervals between plates. Neurons were incubated with compounds for 2.5 h and fixed with 4% paraformaldehyde (EMS; Hatfield, PA) in PBS (Dutscher; Brumath, France) for 20 min at RT, permeabilized with 0.3% Triton-X (Sigma) in PBS for 10 min at RT, and blocked with 5% normal donkey serum (Jackson ImmunoResearch, Ely, UK) and 0.1% Triton-X in PBS for 1 h at RT. Alternatively, neurons in 384-well plates were blocked with 2.5% BSA (Sigma) and 0.1% Triton-X in PBS, up to 14 days at 4°C. Neurons were washed with PBS at RT between each step.

### Proximity ligation assay (PLA)

All components of PLA (Duolink PLA probes and *in situ* detection reagents) apart from the primary and secondary antibodies were from Sigma. PLA was performed following manufacturer’s instructions with minor modifications.^49, 50^ After protein blocking, neurons were incubated with the following primary antibodies overnight at 4°C: BIN1-99D (mouse monoclonal IgG, 1:200; Merck Millipore), Tau (rabbit polyclonal IgG, 1:500; Dako-Agilent), MAP2 (chicken polyclonal IgG, 1:500; Synaptic Systems; Göttingen, Germany), and GFAP (chicken polyclonal IgG, 1:300; Synaptic Systems). Samples were washed with a solution of 0.15 M NaCl (Merck Millipore), 0.01 M Tris (Sigma), 0.05% Tween-20 (Sigma), at pH 7.4 (Buffer A), incubated with PLA probes Mouse-minus and Rabbit-plus (secondary antibodies labeled with complementary DNA strands) in Duolink antibody diluent for 1 h at 37°C, and washed with Buffer A. This was followed by the enzymatic ligation of the two DNA strands, provided that they were in close proximity (< 30 nm),^50^ for 30 min at 37°C and another wash with Buffer A. This was followed by the enzymatic rolling-circle amplification of DNA and hybridization of Cy3-labelled oligonucleotides (PLA orange) for 100 min at 37°C. Samples were then washed with a solution of 0.1 M NaCl and 0.2 M Tris, at pH 7.5 (Buffer B). After the PLA process, samples were incubated with the following secondary antibodies for 1 h at RT: AlexaFlour488 donkey-anti-chicken, AlexaFlour488 donkey-anti-mouse, AlexaFlour647 donkey-anti-rabbit, and DyLight405 donkey-anti-chicken (1:500 for coverslips and 1:1000 for 384-well plates; Jackson ImmunoResearch; West Grove, PA). Coverslips were washed with PBS and mounted in glycerol. 384-well plates were washed with PBS and sealed.

PLA in brain slices was performed with additional modifications.^51^ Slices were first permeabilized with 0.3% Triton in PBS for 30 min and blocked with Duolink blocking solution for 2 h at 37°C. Slices were next treated with the IgG blocking reagent overnight at 4°C and with the protein concentrate, according to manufacturer’s instructions (M.O.M. Basic Kit; Vector Laboratories, Burlingame, CA). Primary antibodies BIN1-99D (1:80), Tau (1:200), and α-tubulin (mouse monoclonal, 1:200; clone DM1A; Sigma) were diluted in the Duolink antibody diluent and incubated overnight at 4°C. Sa mples were washed with Buffer A, incubated with PLA probes Mouse-minus and Rabbit-plus in Duolink antibody diluent for 1 h at 37°C, and washed with Buffer A. This was followe d by DNA ligation for 30 min at 37°C and another wash with Buffer A. This was followed by the enzymatic amplification and PLA hybridization for 2 h at 37°C. Samples were then wa shed with Buffer B and 1:5000 Hoechst (H3569, Thermo Scientific). After the PLA process, samples were incubated with the secondary antibodies AlexaFlour488 donkey-anti-mouse and AlexaFlour647 donkey-anti-rabbit (1:200) for 2 h at RT, followed by several washes with Buffer B. To reduce autofluorescence, the brain slices were treated with 0.1% Sudan Black B (Sigma) in 70% ethanol for 15 min. Samples were then washed with Buffer B and mounted in 90% glycerol.

### Image acquisition and analysis

Coverslips were imaged with LSM 710 confocal microscope (Zeiss, Oberkochen, Germany) using a 40× 1.6 NA objective. Images were acquired at zoom 2 in z-stacks of 0.3 μm interval. 10-13 images per condition were acquired for each of the three independent experiments. Images were deconvoluted using AutoQuantX3 Software (Bitplane, Zurich, Switzerland) and analysed with Imaris Software (Bitplane), using the “surfaces” tool for defining PLA spots, Tau network, and BIN1 puncta in three dimensions. Imaris results were analyzed using a custom MATLAB (MathWorks; Natick, MA) code that removes outliers based on ±3 median absolute deviations (MAD).

384-well plates were imaged using IN Cell Analyzer 6000 Cell Imaging System (GE Healthcare; Little Chalfont, UK) equipped with a Nikon 60× 0.95 NA objective and a CMOS camera. 16 images (2,048 × 2,048 pixels) per well were acquired in four channels (DAPI, dsRed, FITC, and Cy5) using appropriate filter sets and with following acquisition parameters: 2×2 binning; bias = 96.9; gain = 1.0 (Fig. S18). Images were analyzed with Columbus image data storage and analysis system (Perkin Elmer; Waltham, MA) with analysis scripts optimized *via* a custom MATLAB code (Fig. S18B). Optimal analysis scripts were determined separately for each plate.

Brain slices were imaged with Axio Scan Z1 (Zeiss) using a 40× 0.95 NA objective. Images were acquired in 12 z-stacks of 1 μm interval. Regions of interest were marked around the hippocampus during acquisition in each of the 3 independent experiments. PLA spots were analyzed with Imaris using the “surfaces” tool. Imaris results were analyzed using MATLAB after removing outliers based on ±3 MAD.

### HCS script optimization and plate validation

Before image transfer, IN Cell image registration and transfer files were manually edited to import images only from control wells to Columbus, thereby generating the so-called control plates for script optimization and plate validation. Analysis scripts consisted of a series of Columbus commands that determine (i) total Tau staining area and (ii) total area of PLA spots within the Tau network, for each well (Fig. S18A). Four optimization parameters were defined: (i) Tau area threshold in terms of standard deviation (SD) of Tau intensity; (ii) sensitivity parameter for PLA spot detection; (iii) background correction parameter for PLA spot detection; and (iv) minimum PLA spot contrast. Analysis scripts were created by assigning distinct values to each optimization parameter. For example, assigning three distinct values per parameter resulted in 3^4^ = 81 combinations; hence the optimization was performed by running Columbus with 81 separate analysis scripts.

Measured values were corrected for spatial bias (horizontal) using the slope of the line that fits the column averages in the control plate based on the least-squares method. Three values typically used in HCS analysis ^52^ were evaluated: (i) strictly standardized mean difference (β factor, 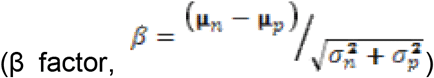); (ii) Z factor 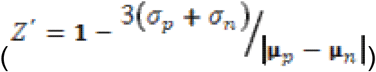; and (iii) signal-to-background ratio 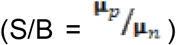, where *μ* and *σ* are mean and standard deviation, and *p* and *n* indicate positive and negative controls. Optimal analysis script was determined as the one with the highest β factor (β ≥ 2), provided that it produced S/B of at least 10. Additional rounds of parameter optimization were performed as deemed necessary.

### Plate analysis and hits selection

Full plates were analyzed with optimal analysis scripts after correcting for local bias in terms of total Tau area, total MAP2 area, and total area of PLA spots within Tau area (Fig. S18C): First the local median of 5x5 wells surrounding the target well calculated and normalized with the plate median excluding edge wells, *i.e.*, *corrected value* = *raw value* / (*local median* / *plate median*). For each plate, compounds affecting network quality, defined as being outside median ± 3 median absolute deviations (MAD) in terms of Tau area or Tau:MAP2 area ratio (edge wells were excluded from these calculations), were excluded (Fig. S19). For each well, corrected PLA:Tau area ratio was normalized by plate mean, excluding edge wells and wells with compounds affecting network quality. After all screenings were performed, mean and SEM of normalized, corrected PLA:Tau area ratio were calculated for each compound, for compounds that did not affect network quality in at least 2 screenings. Compounds potentially affecting BIN1-Tau interaction were determined as those belonging to the top or bottom 5% tiers.

### Validation of selected compounds

Hit validation was performed in a two-step procedure: first, dose-response curves were generated for selected compounds to identify specific effects; second, the impact of selected compounds on BIN1 phosphorylation was assessed through immunoblotting. Since several of the selected compounds had multiple protein targets at 10 μM concentration used in our screen, dose-response experiments were designed to validate the specific effects of the compounds and/or to identify relevant target proteins. Dose-response experiments were performed for 72 selected compounds that induced similar effect on PLA density in all three screens using the same protocol as for the compound screen. Selected compounds were diluted four log scales to obtain a dose-response curve (10nM, 100nM, 1μM and 10μM) and each compound and concentration was tested in three separate plates. Script optimization, plate validation, plate analysis, and well correction and exclusion processes were performed as described above. For each well, corrected PLA:Tau area ratios were normalized by the mean obtained from DMSO-treated wells of the same plate. The means of each compound at 10 μM were compared with the results from screening (conducted at 10 μM), and compounds that had similar effects in both sets of experiments were retained for further analysis. For each compound, dose-response curves were fit with 4-parameter or 3-parameter (where Hill slope is 1) nonlinear regression models, based on the extra sum-of-squares F test using GraphPad Prism 7 (La Jolla, CA). PNC on DIV21 were incubated with selected compounds at 10 μM for 2.5 h and BIN1 and Tau phosphorylation was assessed through immunoblotting.

### Statistical analysis

Statistical analyses were performed in GraphPad Prism 7 or in Matlab. When variables were normally distributed, parametric analyses were applied: one- or two-way analysis of variance (ANOVA), followed by Bonferroni-corrected post hoc tests, Student’s t-test, or one sample t-test. When variables were non-normally distributed, we conducted non-parametric analysis: Kruskal-Wallis ANOVA, followed by Dunn’s test or Wilcoxon signed rank test with Tukey-Kramer correction.

## Acknowledgements

We thank the imaging and animal platforms of the IGBMC and ICS for help, and Nadia Messaddeq, Coralie Spiegelhalter, and Alexia Menuet for technical assistance. We thank the BiCeL platform of the Institut Pasteur de Lille for technical support, and Meryem Tardivel and Antonino Bongiovanni for technical assistance. We thank Hamida Merzougui for technical assistance in recombinant protein preparation. We thank Maxime Verschoore for technical assistance in PLA. We thank Nora L. Salaberry for technical assistance in preparing the summary cartoon. This study was funded by INSERM, CNRS, University of Strasbourg, ANR-10-LABX-0030-INR, a French state fund managed by the ANR under the framework program Investissements d’Avenir (10-IDEX-0002), TGE RMN THC (FR-3050, France), FRABio (University of Lille, CNRS, FR 3688), ANR-BIN-ALZ-15-CE16-0002, PHENOMIN (ANR-10-INBS-07 694), France Alzheimer, the Alzheimer’s Association (BFG-14-318355), the EU Joint Programme – Neurodegenerative Diseases Research (JPND; 3DMiniBrain), Fondation Vaincre Alzheimer (2017 pilot grant), Institut Pasteur de Lille, and Nord-Pas-de-Calais Regional Council. This work was also funded by the Lille Métropole Communauté Urbaine, the French government’s LABEX DISTALZ program (development of innovative strategies for a transdisciplinary approach to Alzheimer’s disease) The NMR facilities were funded by the Nord Regional Council, CNRS, Institute Pasteur de Lille, European Community (FEDER), French Research Ministry and the University of Lille. This work was also funded by the Academy of Finland (307866), Sigrid Jusélius Foundation, and the Strategic Neuroscience Funding of the University of Eastern Finland. M.S. was a fellow of France Alzheimer and received a fellowship from the Fond Paul Mandel de l’Université de Strasbourg. T.M. was supported by a CIFRE fellowship (ANRT/Sanofi), by a Sanofi grant for laboratory supplies, and by the LABEX DISTALZ program.

## Author contributions

I.L., D.K., Y.H., J.L., and J.-C.L. designed and/or supervised research. M.S. and D.M. performed the genotyping and first cohort selection and/or the behavioral experiments in animal model. M.S. performed Tau inclusion labeling and quantifications on mouse brains. S.D. performed PLA and quantification in mouse brains. T.M., S.D., N.M., J.C., A.F., and A.-C.V. performed the *in vitro* experiments. T.M., A.H., F.L., B.D., L.P., D.K., and J.-C.L. performed and/or analyzed the HCS experiments. P.M., M.M., and M.H. performed Western blots and/or analyzed the human sample cohort. A.L., I.M., F.-X.C., and I.L. performed and/or analyzed the NMR experiments. M.S., T.M., P.A., L.P., I.L., D.K., Y.H., J.L., and J.-C.L. wrote and/or revised the paper.

## References

1. Prince, M., Bryce, R., Albanese, E. et al. The global prevalence of dementia: a systematic review and metaanalysis. Alzheimers Dement 9, 63–75.e62 (2013).

2. Zempel, H. & Mandelkow, E. Lost after translation: missorting of Tau protein and consequences for Alzheimer disease. Trends Neurosci 37, 721–732 (2014).

3. Hardy, J. & Selkoe, D.J. The amyloid hypothesis of Alzheimer’s disease: progress and problems on the road to therapeutics. Science 297, 353–356 (2002).

4. Gatz, M., Reynolds, C.A., Fratiglioni, L. et al. Role of genes and environments for explaining Alzheimer disease. Arch Gen Psychiatry 63, 168–174 (2006).

5. Hollingworth, P., Harold, D., Sims, R. et al. Common variants at ABCA7, MS4A6A/MS4A4E, EPHA1, CD33 and CD2AP are associated with Alzheimer’s disease. Nat Genet 43, 429–435 (2011).

6. Lambert, J.C., Heath, S., Even, G. et al. Genome-wide association study identifies variants at CLU and CR1 associated with Alzheimer’s disease. Nat Genet 41, 1094–1099 (2009).

7. Lambert, J.C., Ibrahim-Verbaas, C.A., Harold, D. et al. Meta-analysis of 74,046 individuals identifies 11 new susceptibility loci for Alzheimer’s disease. Nat Genet 45, 1452–1458 (2013).

8. Sims, R., van der Lee, S.J., Naj, A.C. et al. Rare coding variants in PLCG2, ABI3, and TREM2 implicate microglial-mediated innate immunity in Alzheimer’s disease. Nat Genet 49, 1373–1384 (2017).

9. Lambert, J.C. & Amouyel, P. Deciphering genetic susceptibility to frontotemporal lobar dementia. Nat Genet 42, 189–190 (2010).

10. Dourlen, P., Fernandez-Gomez, F.J., Dupont, C. et al. Functional screening of Alzheimer risk loci identifies PTK2B as an in vivo modulator and early marker of Tau pathology. Mol Psychiatry 22, 874–883 (2017).

11. Shulman, J.M., Chipendo, P., Chibnik, L.B. et al. Functional screening of Alzheimer pathology genome-wide association signals in Drosophila. Am J Hum Genet 88, 232–238 (2011).

12. Shulman, J.M., Imboywa, S., Giagtzoglou, N. et al. Functional screening in Drosophila identifies Alzheimer’s disease susceptibility genes and implicates Tau-mediated mechanisms. Hum Mol Genet 23, 870–877 (2014).

13. Beecham, G.W., Hamilton, K., Naj, A.C. et al. Genome-wide association meta-analysis of neuropathologic features of Alzheimer’s disease and related dementias. PLoS Genet 10, e1004606 (2014).

14. Cruchaga, C., Kauwe, J.S., Harari, O. et al. GWAS of cerebrospinal fluid tau levels identifies risk variants for Alzheimer’s disease. Neuron 78, 256–268 (2013).

15. Nelson, P.T., Alafuzoff, I., Bigio, E.H. et al. Correlation of Alzheimer disease neuropathologic changes with cognitive status: a review of the literature. J Neuropathol Exp Neurol 71, 362–381 (2012).

16. Huber, C.M., Yee, C., May, T., Dhanala, A. & Mitchell, C.S. Cognitive Decline in Preclinical Alzheimer’s Disease: Amyloid-Beta versus Tauopathy. J Alzheimers Dis 61, 265–281 (2018).

17. Chapuis, J., Hansmannel, F., Gistelinck, M. et al. Increased expression of BIN1 mediates Alzheimer genetic risk by modulating tau pathology. Mol Psychiatry 18, 1225–1234 (2013).

18. Prokic, I., Cowling, B.S. & Laporte, J. Amphiphysin 2 (BIN1) in physiology and diseases. J Mol Med (Berl) 92, 453–463 (2014).

19. Butler, M.H., David, C., Ochoa, G.C. et al. Amphiphysin II (SH3P9; BIN1), a member of the amphiphysin/Rvs family, is concentrated in the cortical cytomatrix of axon initial segments and nodes of ranvier in brain and around T tubules in skeletal muscle. J Cell Biol 137, 1355–1367 (1997).

20. Ramjaun, A.R., Micheva, K.D., Bouchelet, I. & McPherson, P.S. Identification and characterization of a nerve terminal-enriched amphiphysin isoform. J Biol Chem 272, 16700–16706 (1997).

21. Daudin, R., Marechal, D., Wang, Q. et al. BIN1 genetic risk factor for Alzheimer is sufficient to induce early structural tract alterations in entorhinal cortex-dentate gyrus pathway and related hippocampal multi-scale impairments. bioRxiv (2018).

22. De Rossi, P., Buggia-Prevot, V., Clayton, B.L. et al. Predominant expression of Alzheimer’s disease-associated BIN1 in mature oligodendrocytes and localization to white matter tracts. Mol Neurodegener 11, 59 (2016).

23. Miyagawa, T., Ebinuma, I., Morohashi, Y. et al. BIN1 regulates BACE1 intracellular trafficking and amyloid-beta production. Hum Mol Genet 25, 2948–2958 (2016).

24. McKenzie, A.T., Moyon, S., Wang, M. et al. Multiscale network modeling of oligodendrocytes reveals molecular components of myelin dysregulation in Alzheimer’s disease. Mol Neurodegener 12, 82 (2017).

25. Calafate, S., Flavin, W., Verstreken, P. & Moechars, D. Loss of Bin1 Promotes the Propagation of Tau Pathology. Cell Rep 17, 931–940 (2016).

26. Malki, I., Cantrelle, F.X., Sottejeau, Y. et al. Regulation of the interaction between the neuronal BIN1 isoform 1 and Tau proteins - role of the SH3 domain. Febs j 284, 3218–3229 (2017).

27. Polydoro, M., Acker, C.M., Duff, K., Castillo, P.E. & Davies, P. Age-dependent impairment of cognitive and synaptic function in the htau mouse model of tau pathology. J Neurosci 29, 10741–10749 (2009).

28. Andorfer, C., Kress, Y., Espinoza, M. et al. Hyperphosphorylation and aggregation of tau in mice expressing normal human tau isoforms. J Neurochem 86, 582–590 (2003).

29. Andorfer, C., Acker, C.M., Kress, Y. et al. Cell-cycle reentry and cell death in transgenic mice expressing nonmutant human tau isoforms. J Neurosci 25, 5446–5454 (2005).

30. Buee, L., Bussiere, T., Buee-Scherrer, V., Delacourte, A. & Hof, P.R. Tau protein isoforms, phosphorylation and role in neurodegenerative disorders. Brain Res Brain Res Rev 33, 95–130 (2000).

31. Sottejeau, Y., Bretteville, A., Cantrelle, F.X. et al. Tau phosphorylation regulates the interaction between BIN1’s SH3 domain and Tau’s proline-rich domain. Acta Neuropathol Commun 3, 58 (2015).

32. Bauerfeind, R., Takei, K. & De Camilli, P. Amphiphysin I is associated with coated endocytic intermediates and undergoes stimulation-dependent dephosphorylation in nerve terminals. J Biol Chem 272, 30984–30992 (1997).

33. Floyd, S.R., Porro, E.B., Slepnev, V.I. et al. Amphiphysin 1 binds the cyclin-dependent kinase (cdk) 5 regulatory subunit p35 and is phosphorylated by cdk5 and cdc2. J Biol Chem 276, 8104–8110 (2001).

34. Lasorsa, A., Malki, I., Cantrelle, F.X. et al. Structural basis of Tau interaction with BIN1 and regulation by Tau phosphorylation. Front Mol Neurosci 11, 421.

35. Qi, H., Prabakaran, S., Cantrelle, F.X. et al. Characterization of Neuronal Tau Protein as a Target of Extracellular Signal-regulated Kinase. J Biol Chem 291, 7742–7753 (2016).

36. Broadbent, N.J., Squire, L.R. & Clark, R.E. Spatial memory, recognition memory, and the hippocampus. Proc Natl Acad Sci U S A 101, 14515–14520 (2004).

37. Van Cauter, T., Camon, J., Alvernhe, A. et al. Distinct roles of medial and lateral entorhinal cortex in spatial cognition. Cereb Cortex 23, 451–459 (2013).

38. Chapuis, J., Flaig, A., Grenier-Boley, B. et al. Genome-wide, high-content siRNA screening identifies the Alzheimer’s genetic risk factor FERMT2 as a major modulator of APP metabolism. Acta Neuropathol 133, 955–966 (2017).

39. Glennon, E.B., Whitehouse, I.J., Miners, J.S. et al. BIN1 is decreased in sporadic but not familial Alzheimer’s disease or in aging. PLoS One 8, e78806 (2013).

40. Holler, C.J., Davis, P.R., Beckett, T.L. et al. Bridging integrator 1 (BIN1) protein expression increases in the Alzheimer’s disease brain and correlates with neurofibrillary tangle pathology. J Alzheimers Dis 42, 1221–1227 (2014).

41. Ferrer, I. Oligodendrogliopathy in neurodegenerative diseases with abnormal protein aggregates: The forgotten partner. Prog Neurobiol 169, 24–54 (2018).

42. Tucker, K.L., Meyer, M. & Barde, Y.A. Neurotrophins are required for nerve growth during development. Nat Neurosci 4, 29–37 (2001).

43. Beaudoin, G.M., 3rd, Lee, S.H., Singh, D. et al. Culturing pyramidal neurons from the early postnatal mouse hippocampus and cortex. Nat Protoc 7, 1741–1754 (2012).

44. Kaech, S. & Banker, G. Culturing hippocampal neurons. Nat Protoc 1, 2406–2415 (2006).

45. Long, K., Mohan, C., Anderl, J. et al. Analysis of autophagosome formation using lentiviral biosensors for live fluorescent cellular imaging. Methods Mol Biol 1219, 157–169 (2015).

46. Braak, H., Alafuzoff, I., Arzberger, T., Kretzschmar, H. & Del Tredici, K. Staging of Alzheimer disease-associated neurofibrillary pathology using paraffin sections and immunocytochemistry. Acta Neuropathol 112, 389–404 (2006).

47. Natunen, T., Parrado, A.R., Helisalmi, S. et al. Elucidation of the BACE1 regulating factor GGA3 in Alzheimer’s disease. J Alzheimers Dis 37, 217–232 (2013).

48. Welburn, J. & Endicott, J. Methods for preparation of proteins and protein complexes that regulate the eukaryotic cell cycle for structural studies. Methods Mol Biol 296, 219–235 (2005).

49. Bagchi, S., Fredriksson, R. & Wallen-Mackenzie, A. In Situ Proximity Ligation Assay (PLA). Methods Mol Biol 1318, 149–159 (2015).

50. Soderberg, O., Leuchowius, K.J., Gullberg, M. et al. Characterizing proteins and their interactions in cells and tissues using the in situ proximity ligation assay. Methods 45, 227–232 (2008).

51. Gomes, I., Sierra, S. & Devi, L.A. Detection of Receptor Heteromerization Using In Situ Proximity Ligation Assay. Curr Protoc Pharmacol 75, 2.16.11-12.16.31 (2016).

52. Bray, M.A., Carpenter, A., Imaging Platform, B.I.o.M.I.T. & Harvard in Assay Guidance Manual. (eds. G.S. Sittampalam et al.) (Eli Lilly & Company and the National Center for Advancing Translational Sciences, Bethesda (MD); 2004).

